# Evolution and genomic signatures of spontaneous somatic mutation in *Drosophila* intestinal stem cells

**DOI:** 10.1101/2020.07.20.188979

**Authors:** Nick Riddiford, Katarzyna Siudeja, Marius van den Beek, Benjamin Boumard, Allison J. Bardin

## Abstract

Spontaneous mutations can alter tissue dynamics and lead to cancer initiation. While large-scale sequencing projects have illustrated processes that influence somatic mutation and subsequent tumour evolution, the mutational dynamics operating in the very early stages of cancer development are currently not well understood. In order to explore mutational dynamics in the early stages of cancer evolution we exploited neoplasia arising spontaneously in the *Drosophila* intestine. We analysed whole-genome sequencing data through the development of a dedicated bioinformatic pipeline to detect structural variants, single nucleotide variants, and indels. We found neoplasia formation to be driven largely through the inactivation of *Notch* by structural variants, many of which involve highly complex genomic rearrangements. Strikingly, the genome-wide mutational burden of neoplasia - at six weeks of age - was found to be similar to that of several human cancers. Finally, we identified genomic features associated with spontaneous mutation and defined the evolutionary dynamics and mutational landscape operating within intestinal neoplasia over the short lifespan of the adult fly. Our findings provide unique insight into mutational dynamics operating over a short time scale in the genetic model system, *Drosophila melanogaster*.

## Introduction

The accumulation of mutations in somatic tissues plays a major role in cancer and is proposed to contribute to ageing (Al Zouabi and Bardin 2020). While the majority of mutations acquired throughout life are harmless, some alter cellular fitness and become subject to the selective forces operating on cells and tissues. Mutations that confer a selective advantage can lead to the formation of a clonal population of mutant cells under positive selection. Such events, termed driver mutations, underscore cancer formation and, as such, have been the subject of extensive investigation (ICGC/TCGA Pan-Cancer Analysis of Whole Genomes Consortium 2020; Rheinbay et al. 2020; Alexandrov et al. 2020; Bailey et al. 2018). Importantly, these initiating mutations are thought to arise in normal cells, and can therefore provide key insights into the mutational processes operative in pre-cancerous states. Large-scale sequencing projects have detailed the mutational burdens of human cancer genomes and have revealed the repertoire of somatic mutations driving cancer formation, illuminating the biological processes underlying somatic mutation. Cancer genomes, however, represent the end-point of a long evolutionary process that shapes the mutational landscape of tumours. Similarly, the mutations recently described to arise in aged normal cells and early-stage cancers represent the result of many years of selective pressure and mutational dynamics (Martincorena et al. 2015, 2018; Lee-Six et al. 2019; Moore et al. 2020; Yokoyama et al. 2019). Knowledge of mutational processes operative in the very earliest stages of cancer is therefore currently incomplete.

Our previous work has established the Drosophila midgut as an excellent model system for understanding somatic mutation in an adult tissue-specific stem cell population (Siudeja et al. 2015). In this tissue, intestinal stem cells (ISCs) self-renew and divide to give rise to two differentiated cell types: absorptive enterocytes (ECs) and secretory enteroendocrine cells (EEs) (Micchelli and Perrimon 2006; Ohlstein and Spradling 2006). We have previously shown that during ageing, 12% of wild-type male flies harbour spontaneous mutations that inactivate the X-linked tumour-suppressor gene *Notch*, driving hyperproliferation of ISCs and EEs and resulting in neoplasm formation (Siudeja et al. 2015).

Here, we take advantage of the spontaneous formation of neoplasia in the intestine of the fruit fly to investigate the processes underlying early somatic mutation and evolution within a clonal cell population. We analysed whole-genome sequencing data of neoplasia to characterise structural variants, single nucleotide variants (SNVs), and small insertions/deletions (indels) occurring genome-wide in ISCs. We found that inactivation of *Notch*, in normal stem cells, is driven by structural variation, many of which are complex rearrangements likely driven by replicative processes. Moreover, we detect transposable element (TE) sequences associated with structural variants. Exploiting the clonal nature of the neoplasia, we categorised mutational timing relative to the driver mutations in *Notch*. Our data suggest that over a period of weeks, SNVs are acquired more rapidly than structural variants and indels. Strikingly, in neoplasia from six-week-old male flies, we detected a genome-wide mutational burden similar to that found in several human cancers (Alexandrov et al. 2013a). Our experimental setup exploits both the small genome size and short lifespan of *Drosophila* to allow us to rapidly assay the full spectrum of mutations across the genome in aged stem cells. This provides us with a unique opportunity to investigate both somatic mutations in normal stem cells, as well as those occurring in the very early stages of neoplasm development.

## Results

### 1. A comprehensive pipeline to detect somatic structural variation in *Drosophila* intestinal stem cells

We have previously shown that ISCs of the *Drosophila* midgut spontaneously acquire structural variants during ageing that disrupt tissue homeostasis via the inactivation of the X-linked tumour suppressor gene *Notch* (Siudeja et al. 2015). In males, a single inactivation of *Notch* in ISCs leads to neoplasm formation, comprising a highly proliferative and rapidly expanding clonal population of *Notch* mutant ISCs and EE cells. Here, we leverage this system to dissect the mechanisms underlying *Notch* inactivation, and characterise the landscape of somatic mutations in ageing stem cell genomes via the development of a robust bioinformatic pipeline.

In order to define spontaneously arising somatic mutations, we analysed whole-genome sequencing data generated from 35 intestinal neoplasia. To enable us to discern somatic events, we compared neoplasia with the head from the same fly as a direct control and, consistent with human cancer studies, we will refer to sequenced neoplasia and heads as “tumour” and “normal” samples, respectively. Three samples were re-analysed from a previously published dataset (Siudeja et al. 2015), and the remaining samples are described in further detail in (Siudeja et al. 2020). Using this approach, we can exploit the clonal nature of the tumours to identify somatic mutations in ISCs that are difficult to detect in genetically mosaic adult tissues (Fig. 1a). To accurately characterise the full spectrum of structural variants, we developed a pipeline that combines multiple best-practise approaches and applies stringent filters with several novel annotation method. This pipeline incorporates readdepth-based approaches for detecting copy number variants (CNVs), as well those utilising read-mapping signatures. As a part of the pipeline we developed several novel tools to filter and annotate structural variant calls (Fig. 1b; Fig. S1a; Methods; Supplementary Methods). To ensure that only somatic variants were considered, we constructed a panel of normals (PON) by combining all normal samples, and filtered out variants that were found in any of these samples. Orthogonal to this current study, we have developed a bioinformatic tool to tag paired-end reads that partially map to, or have mates that map to, non-reference DNA sequences (Methods; (Siudeja et al. 2020)). Here, we utilised this tool to tag reads associated with TEs as well as enteric bacterial and viral species. In doing so, we were able to filter out microbial genomic sequences prevalent in the gut samples that artificially map to the *D. melanogaster* genome, as well as identify germline TEs that are not present in the reference genome.

**Fig. 1.**
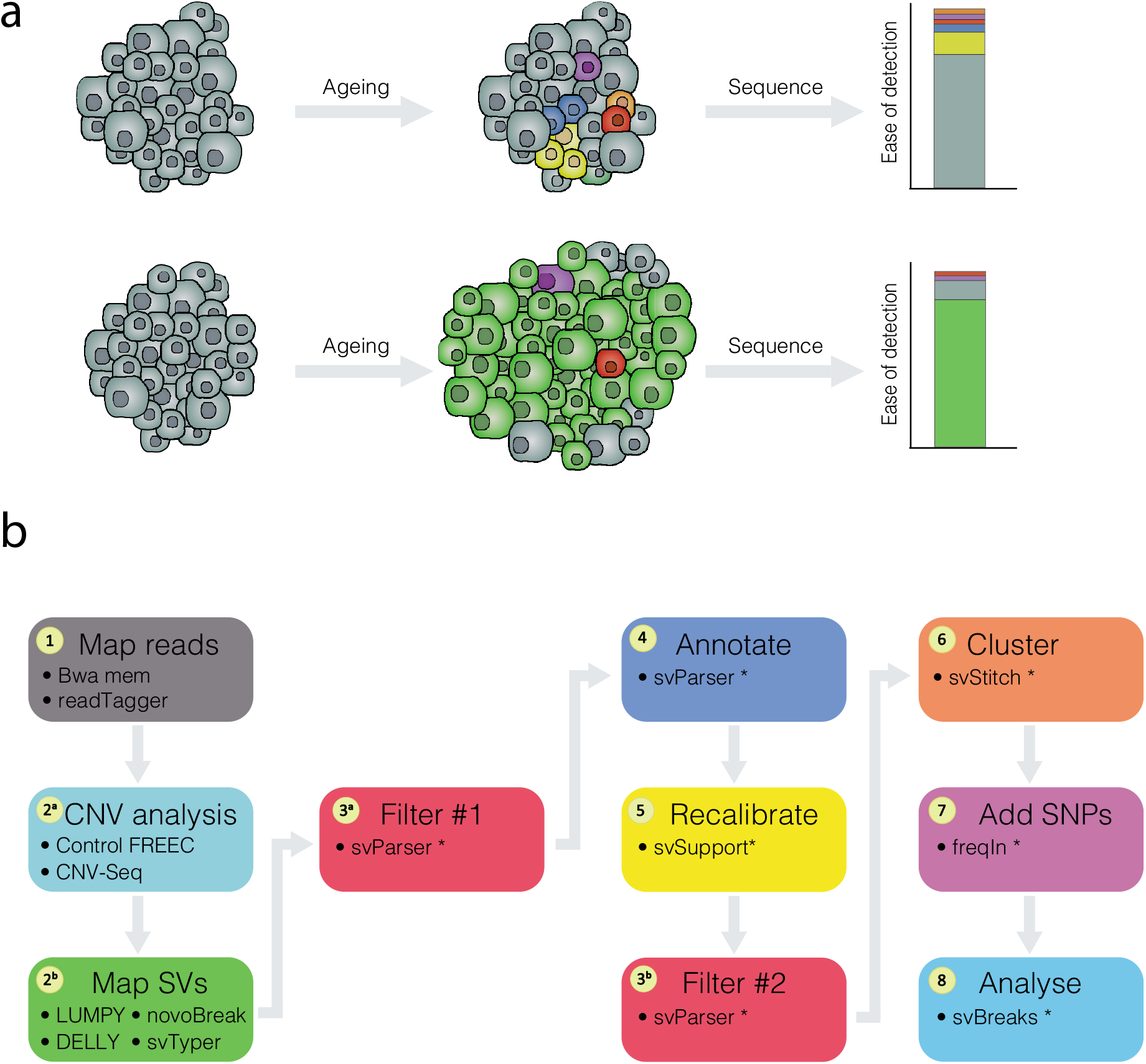
Clonal expansion of ISCs can be exploited to detect somatic mutations. (**a**) During ageing, normal cells (grey) acquire somatic mutations (coloured), typically restricted to small populations of cells, that give rise to genetic mosaicism within the tissue. Bulk DNA-sequencing of such tissues fails to detect somatic mutations, as they are present in such small numbers. Somatic mutations occurring in an ISC (green) are inherited by the cell’s progeny, and in the context of a neoplasm, are present in many cells within the tissue. As a result, sequencing of neoplasia increases the ability to detect somatic mutations in wild-type tissue. (**b**) A comprehensive bioinformatic pipeline was created in order to accurately detect and characterise structural variants from sequenced neoplasia. We have developed multiple packages to enable us to tag reads that map to multiple genomes (Siudeja et al. 2020), and filter and annotate structural variant breakpoints (svParser, svSupport, freqIn; Methods; Supplementary Methods). Our pipeline utilises multiple approaches to detect structural variants, and applies stringent filtering steps before annotating variants. Steps marked by an asterisk indicate bioinformatic tools developed for this study.

In cases where multiple structural variant breakpoints were found within small (5 kb) windows, individual calls were collapsed into unified “complex” events and recorded as a distinct class. The bioinformatic pipeline that we have developed therefore enables the comprehensive detection of multiple types of somatic mutation, combining stringent filters with extensive annotation approaches to investigate somatic mutation in stem cell-derived neoplasia of the *Drosophila* intestine.

### 2. Diverse mutational events inactivate *Notch* in normal ISCs

In male flies, a single loss of *Notch* in ISCs is sufficient to drive tumour formation (Siudeja et al. 2015), providing a useful model locus to understand mechanisms of tumour suppressor inactivation. These mutations occur in normal ISCs, and as such allow us to characterise somatic mutations in a pre-neoplastic state. We therefore initially focussed on mutations affecting *Notch*.

In the 35 tumour samples analysed, we found *Notch* to be inactivated via multiple different classes of structural variants with lengths ranging from 200 bps to 550 kb (Fig. 2a). These included deletions: (20/35; 57.1%), complex rearrangements (8/35; 22.8%); an inversion (1/35, 2.9%) and one translocation (1/35, 2.9%). In five samples (P15, P37, P47, P51, D5), we found no evidence for inactivation of *Notch* by a structural variant. However, extending our search to other Notch pathway components revealed that sample P37 had multiple structural variants spanning 46 kb, which we hypothesise resulted in the biallelic inactivation of *kuzbanian*, a protease required for *Notch* activation, likely responsible for tumour formation. While a further investigation of somatic TE insertions in this system will be described elsewhere ((Siudeja et al. 2020)), we note that the remaining four samples for which we did not find support for a structural variant in *Notch* (P15, P47, P51, D5), had evidence supporting *de novo* transposable element insertion in *Notch*, likely causing its inactivation.

**Fig. 2.**
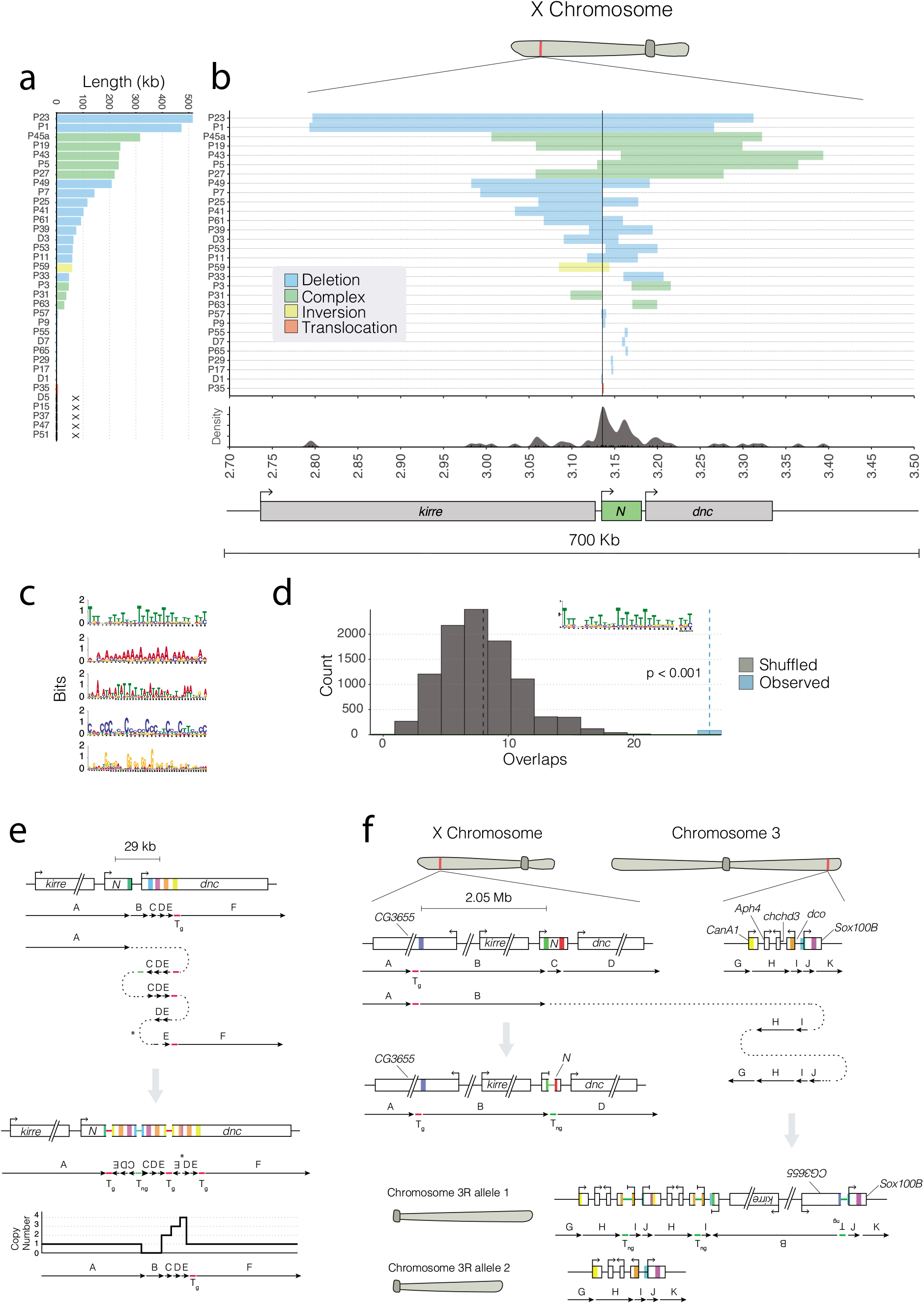
*Notch* is inactivated by multiple different mutational events. (**a**) Structural variants affecting *Notch* in each sample vary in size and class. Across all samples, we find *Notch* to be inactivated by deletions (20/35; blue), complex rearrangements (8/35; green), an inversion (1/35, sample P59; yellow) and one translocation (1/35, sample P35; red). In four samples (marked as ‘X’) we did not detect a structural variant in *Notch*. (**b**) The distribution of structural variant breakpoints over the *Notch* locus. While we did not detect mutational hotspots in *Notch*, we observed a clustering (density plot) of breakpoints around the TSS of *Notch*, indicated by a black vertical line. (**c**) Motifs found enriched +/- 500 bps of *Notch* breakpoint are highly repetitive. Permutation tests show that breakpoint flanking sequences are significantly enriched (p < 0.001) for poly(dA:dT) sequences. We observed 25 overlaps between breakpoint flanking sequences and poly(dA:dT) sequences (blue vertical dashed line), and in 10,000 permutations we detected a median of 7 overlaps (black vertical dashed line). (**e, f**) Putative explanations of two complex genomic rearrangements inactivating *Notch* based on read support and CNV calls. Schematics show genomic regions before (top) and after (bottom) each rearrangement. Coloured boxes represent breakpoints, with the resulting genomic adjacencies shown below. Arrows indicate the order and orientation of genomic regions, and dotted lines represent a template switching event during DNA replication. Transposable elements are shown as genomic regions, with non-germline sequences (T_ng_) shown in green, and germline sequences (T_g_) shown in red. (**e**) A complex event generating a deletion in region B, followed by an inverted quadruplication of downstream sequence (regions C, D and E), flanked by TE sequences. We detected a 12 bps locally templated insertion (indicated by an asterisk) at the breakpoint junction between regions C and D. A schematic of the resulting copy number profile is shown below. (**f**) A translocation from *Notch* to chromosome 3R. A 2.05 Mb region upstream of *Notch* (region B) is incorporated onto chromosome 3R and the entire region is subsequently duplicated. The region immediately upstream of the translocation breakpoint on the X chromosome (region C) is deleted, and we detect TE sequence at the breakpoint, as well as at the 5’ breakpoint of region J, and the junction between regions H and I. In this model, one copy of chromosome 3R contains the rearranged region from the X chromosome (labelled allele 1), while the other is unaltered (allele 2).

In human cancer genomes, structural variant breakpoints are distributed non-uniformly, and are commonly found to be located in regions of the genome that are inherently prone to double-strand break (DSB) damage (Glodzik et al. 2017). In order to establish whether hotspot regions existed in the *Notch* locus, we examined the distribution of breakpoints (Fig. 2b; Fig. S2a). While no two breakpoints had the same genomic position, we observed clusters of breakpoints in close proximity (+/- 5 kb; Fig. 2b), including close to the transcription start site (TSS) of *Notch* (breakpoints in 7/35 samples within +/- 2 kb of the TSS; Fig. S2a).

We next investigated whether breakpoints in *Notch* shared underlying sequence similarity that could provide insight into the mechanisms involved in their formation. In particular, we searched for sequences that have the potential to form alternative DNA conformations (non-B-form DNA), including cruciform DNA, short inverted repeats (SIR) and G-quadruplexes, all of which can promote genome instability (Lu et al. 2015; Kurahashi et al. 2004; Paeschke et al. 2011). We extracted the sequence +/- 500 bps from each breakpoint in *Notch* and performed permutation tests on the overlap between repeats and these breakpoint-flanking regions. We did not find significant enrichment of inverted repeats and G-quadruplexes around we used MEME (Bailey and Elkan 1994) to perform *de novo* motif discovery on breakpoint regions. All of motifs recovered were highly repetitive, comprising mono- or di-nucleotide repeats, that resembled microsatellites - tandem repeats of 1-6 bp, sequences which have been previously shown to be prone to mutation due to replication slippage, mismatch repair or fork-stalling during DNA replication (Fig. 2c; (Gadgil et al. 2017)). breakpoints. To determine whether other sequences might be associated with breakpoints,

Of particular interest, the two most highly overrepresented motifs we found - mononucleotide A/T repeats - were strikingly similar to the poly(dA:dT) tracts recently identified as preferential sites of replication fork collapse upon induction of replication stress by hydroxyurea (Tubbs et al. 2018). In light of this association between poly(dA:dT) tracts and replication fork collapse, we then performed a genome-wide search for these motifs, before performing permutation tests on the overlap between motif occurrences and the regions flanking *Notch* breakpoints. This analysis revealed that breakpoint regions were significantly enriched for poly(dA:dT) tracts when compared to the genomic region surrounding *Notch* (Fig. 2d), with 26/42 (62%) breakpoint regions containing one or more poly(dA:dT) tract. The high enrichment of poly(dA:dT) tracts at breakpoints supports the hypothesis that replication fork collapse may promote many of the structural variants observed in *Notch*.

Interestingly, replication fork collapse has been proposed to generate complex structural variants through a mechanism known as microhomology-mediated break-induced replication (MMBIR), related to the previously proposed fork stalling and template switching (FoSTeS) (Lee et al. 2007; Hastings et al. 2009; Carvalho and Lupski 2016; Li et al. 2020). Interestingly, a quarter of the structural variants inactivating *Notch* comprised complex rearrangements (8/35; 25.7%) that are difficult to explain via classical models of DSB repair such as non-homologous end-joining (NHEJ) or homologous recombination (HR) pathways, which would require multiple independent events. Replication-driven rearrangements are often associated with short stretches of sequence homology (microhomology) as well as insertions at breakpoint junctions. Indeed, many of the complex events in *Notch* harboured locally templated insertions at breakpoint junctions, indicative of being generated by replicative mechanisms such as MMBIR ((Carvalho et al. 2013; Liu et al. 2012); Supplementary Table 1). Below, we present two complex events that we hypothesise to have been driven by replicative mechanisms, based on read support and CNV calls.

In sample P63, a 23 kb deletion with breakpoints in the last exon of *Notch* and the second intron of *dunce* was followed by a 1 kb duplication, a 4 kb inverted triplication and a 1 kb quadruplication (Fig. 2e). On inspecting breakpoint junctions, we detected reads mapping to an I-element TE at both the 5’ breakpoint of the deletion and the 5’ breakpoint of the quadruplication, as well as a short, inserted, sequence at the breakpoint junction of the inverted region that was likely locally templated. This sort of highly complex rearrangement, including copy number (CN) changes, inverted regions and templated insertions is likely best explained by template switching of a replicative polymerase during replication (Carvalho et al. 2013; Liu et al. 2012).

In another sample (P35), we found *Notch* to have been inactivated by a translocation with breakpoints located in the 1st intron of *Notch* and 2 kb upstream of *Sox100B* on 3R (Fig. 2f). Interestingly, we found the breakpoint on 3R to be located within a tandem duplication. By inspecting read depth on the X, we detected a CNV region 2.05 Mb upstream of the translocation breakpoint in *Notch*, with a putative 5’ breakpoint in a germline I-element TE. The region immediately downstream of the translocation breakpoint was deleted (575 bps), and we detected I-element mapping reads at this breakpoint junction, as well as at several junctions on 3R. Considering the tandem duplication on 3R, one explanation for such a configuration is that the entire 2.05 Mb region upstream of *Notch* was incorporated on chromosome 3R, which was then subsequently duplicated as part of a complex rearrangement (Fig. 2f).

Taken together with the enrichment of poly(dA:dT) at breakpoints, such signatures of template switching in complex events suggests that replication fork collapse may underlie inactivation of *Notch*.

### 3. Transposable elements and viral inserts at structural variant breakpoints in *Notch*

Considering the presence of TE-tagged reads at the breakpoints of several complex events, we extended our breakpoint characterisation to scrutinise reads tagged as either mapping to, or having a mate mapping to, TE sequences (Methods). We frequently observed evidence of TE sequences at breakpoints within the *Notch* locus (11/30 *Notch-*inactivating structural variants). Of these, 8/11 were at deletion breakpoints, representing 40% of the deletions within *Notch*, and 3/11 were at breakpoints of complex rearrangements, comprising 37.5% of the complex variants affecting *Notch*. The frequent association of TE sequences with deletions and complex variants, motivated us to further investigate the potential mechanisms underlying their formation.

To explore TE-involvement in structural variant formation, we characterised TEs at breakpoints according to their family and somatic status (somatic or germline) and identified four classes of event, suggestive of distinct mechanisms (Fig. 3a-d). In Class I events (3/11), TE-mapping reads were detected at both breakpoints, with no supporting evidence for TE sequences in the corresponding normal tissue. We believe that this represents the integration of either a full length TE, or TE fragment, at breakpoint junctions (Fig. 3a). In Class II events (3/11) we observed that both breakpoints were located within germline TEs. Here, we suspect that variants were generated via non-allelic homologous recombination (NAHR) between two germline TEs with high sequence similarity (Fig. 3b) as previously reported (Robberecht et al. 2013).

**Fig. 3.**
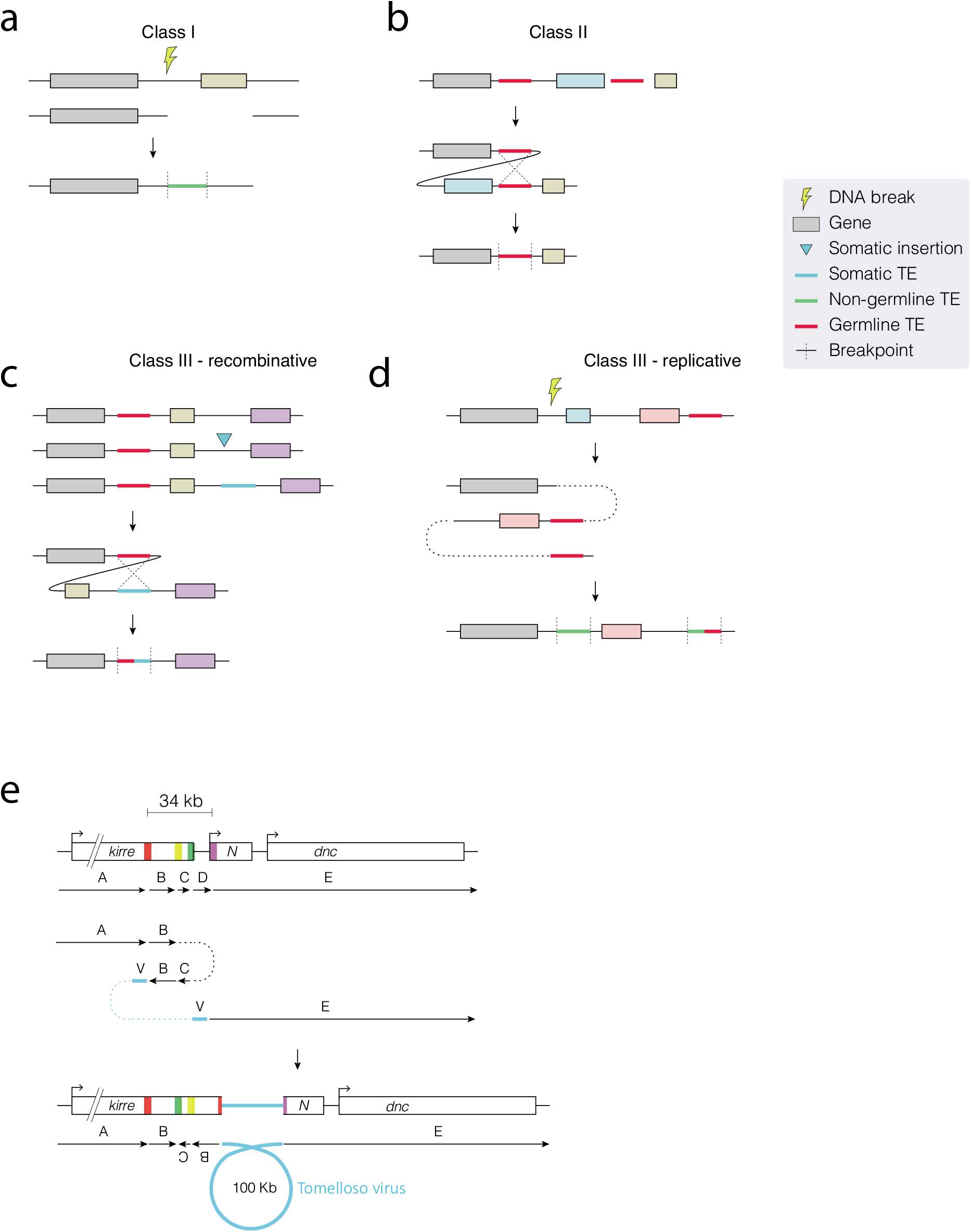
Transposable element sequences and viral insertions in *Notch*-inactivating structural variants. (**a-d**) Schematics show putative mechanisms of rearrangement that could explain the signatures of TE involvement detected in *Notch*-inactivating structural variants. In each class, the uppermost schematic shows a hypothetical genomic region, with genes indicated by coloured boxes to help visualise the resulting rearrangement (shown at the bottom). Germline (Tg; red), non-germline (Tng; green) and somatic (Ts; blue) TE sequences are shown as bold coloured lines, and dotted lines indicate the joining of non-contiguous genomic regions, with vertical dotted lines showing breakpoints in the rearranged sequence. (**a**) In class I events, read evidence supports a TE or TE fragment integrated at the breakpoint junction. We hypothesise that the TE sequence was integrated during DNA repair. (**b**) In class II events, two germline TE sequences are found at breakpoint junctions. We hypothesise that these sequences underwent non-allelic homologous recombination, deleting the central region. (**c, d**) Class III events have evidence for a non-germline TE sequence at one breakpoint junction and germline TE sequence at another. Two interpretations for the breakpoint signatures present in class III events. (**c**) In the first, a recombinative explanation posits that a *de novo* TE sequence is inserted (blue arrow) upstream of a germline TE belonging to the same family. Recombination between the two TEs deletes the central region. (**d**) A second possible explanation of class III breakpoint signatures, wherein DNA damage is repaired by a replicative polymerase that erroneously integrates TE sequence into one or several of the breakpoint junctions. This results in a *de novo* TE signature, and in this example, we illustrate this within the context of an inverted duplication of the genomic region immediately downstream of the initial DSB. (**e**) A schematic showing genomic regions before (top) and after (bottom) the integration of a fragment of a viral genome in the context of a complex rearrangement detected in sample P31. While our sequencing data support this configuration, it is possible that alternative explanations exist. Coloured boxes represent breakpoints, and arrows indicate the order and orientation of genomic regions. Dotted lines represent a template switching event during DNA replication. Blue lines indicate viral DNA sequence.

In Class III events (5/11), one breakpoint originated in a germline TE, while the other breakpoint was mapped to non-germline TE sequence. Here, we classify TE sequences as being “non-germline”, to distinguish them from putative somatic insertions. Importantly, we observed that both germline and non-germline TE sequences belonged to the same family and here, we consider two explanations for such breakpoint signatures. First, given the sequence similarity, it is possible that structural variants were generated as result of a homologous recombination event between a somatic TE and the germline element (Fig. 3c). However, an alternative possibility is that a germline TE acted as a substrate for template switching during DNA replication, copying TE sequence into a novel locus. This would explain the association of this class with the breakpoints of complex rearrangements in *Notch* that likely arose via replicative mechanisms (Fig. 3d). That we detect the involvement of TEs in so many of the structural variants in *Notch* highlights the role that TE sequences may play in influencing somatic mutation, as well as underscoring the complexity of mutations inactivating a model tumour-suppressor locus.

In addition to detecting TE presence at breakpoints, we identified breakpoints at reads whose mates mapped to the double-stranded DNA (dsDNA) nudivirus Tomelloso in one sample (P31; (Palmer et al. 2018). The breakpoint read orientation was consistent with a ~100 kb fragment of viral DNA integrated into the *Drosophila* genome as part of a complex rearrangement (Fig. 3e) and to our knowledge, this is the first known example of a somatic dsDNA viral insertion in *Drosophila*. We did not detect virus-associated variants in other samples or genomic loci, and found no correlation between the number of mutations detected and the viral load per sample (Fig. S3).

### 4. The mutational burden of ISCs

Due to the difficulty of accurately identifying mutations present in small numbers of cells from highly mosaic adult tissues, somatic mutations arising in normal adult tissues are challenging to characterise. Our experimental set-up allows us to investigate such mutations and, as well as understanding the variants affecting *Notch*, we are able to exploit the clonal nature of the ISC-derived tumours to interrogate somatic mutations in adult stem cell genomes.

First, we extended our structural variant analysis to consider the genome-wide distribution and characteristics of all instances of somatic structural variation (Supplementary Table 2 - genome-wide variants). Overall, we find multiple classes of structural variants distributed throughout the mappable genome, with no breakpoint clustering apparent outside of the *Notch* locus (Fig. 4a). In total we found 618 structural variants across all samples (median: 6 per sample), 36% of which (222/618) were translocations where the fraction of supporting reads was low, and were likely to be highly subclonal to the original mutation in *Notch* (Fig. 4b, c; Fig. S4a). Interestingly, the relative frequency of structural variant classes genome-wide was quite distinct from those observed in *Notch*. We found translocations to be enriched genomewide, whereas both deletions and complex rearrangements considerably more frequently observed in *Notch* than genome-wide variants (Fig. 4b). It is likely that this difference is inherent to our assay, which selects for *Notch*-inactivating events. Owing to the greater disruptive potential of variants involving deletion, it is perhaps not surprising that these classes of events are more frequently observed in *Notch* relative to genome-wide variants.

**Fig. 4.**
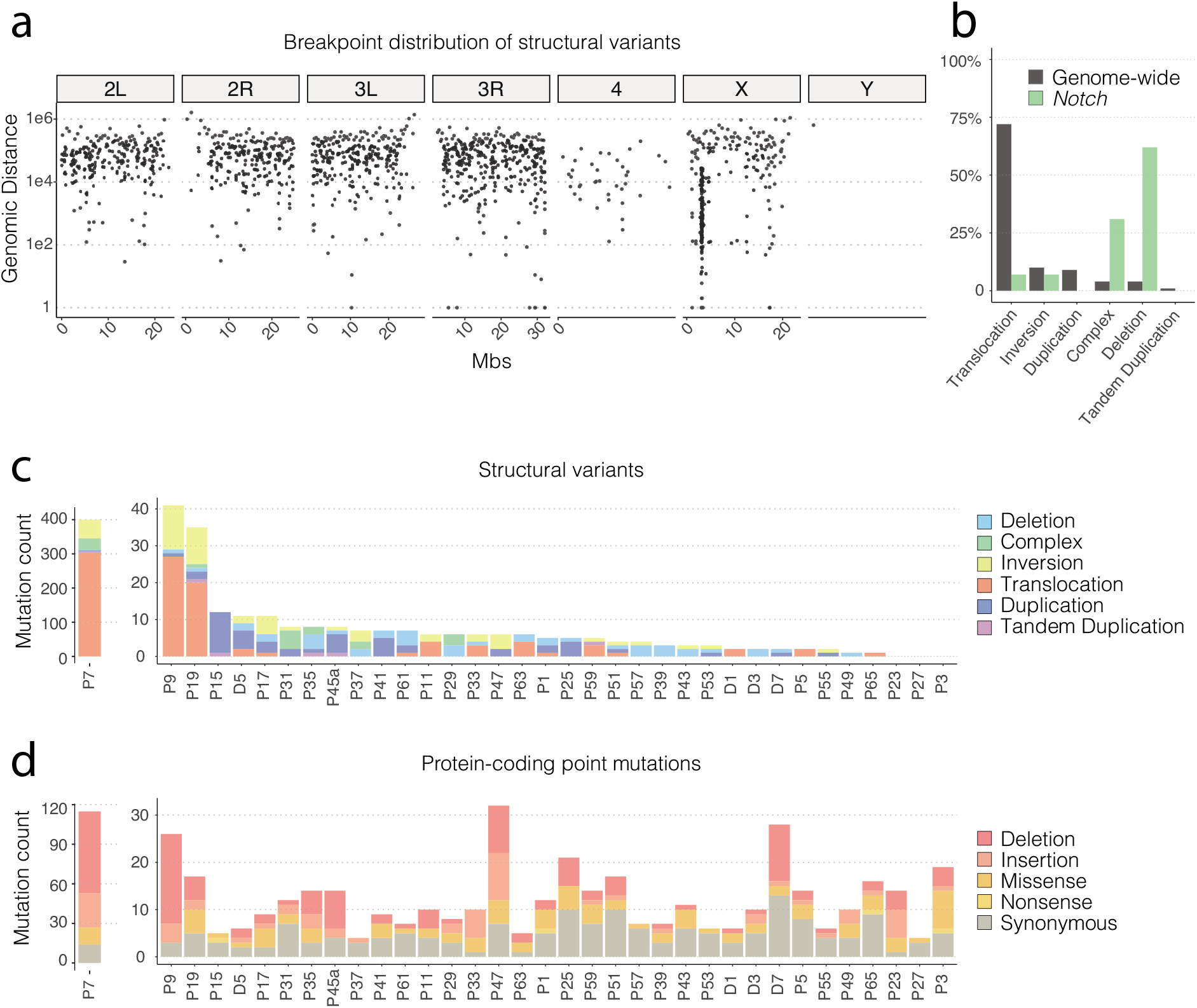
Multiple classes of somatic mutations were detected genome-wide. (**a**) A rainfall plot showing the distances between structural variant breakpoints across the genome. The y axis shows the Log10 distance between consecutive breakpoints, with lower numbers representing smaller distances between breakpoints. (**b**) The percentage contribution of different structural variant classes to the total number of mutations identified genome-wide (grey bars) and in *Notch*-inactivating variants (green bars). (**c, d**) The number of each class of structural variant (**c**) and protein-coding point mutation (**c**; SNVs and indels) observed across samples. In both **c** and **d**, sample P7 is plotted on a separate axis to aid visualisation.

Next, to characterise the full spectrum of somatic mutation in ISC genomes, we extended our analysis to include point mutations, including SNVs and indels. As with the structural variant analysis, SNVs and indels were detected using multiple best-practice approaches, genotyped against a PON, and stringently filtered to ensure a high-quality call set of somatic events (Methods). As well as extensive manual inspection of calls, we also assessed the quality of SNVs by calculating the transition/transversion (Ti/Tv) ratio across samples. Considering that there are more possible transversions (A↔C, A↔T, C↔G, G↔T) than transitions (A↔G or C↔T), the Ti/Tv ratio is often used as a quality control to discriminate non-random substitution rates, generally indicated by Ti/Tv values in excess of 0.5. We find a genome-wide Ti/Tv ratio of 0.9 which, while substantially lower than values reported in mammalian data sets (e.g. (Bainbridge et al. 2011)), is broadly consistent with comparable *Drosophila* datasets (Petrov and Hartl 1999; Keller et al. 2007). This observed difference is largely explained by the lack, or very low levels, of DNA methylation in *Drosophila* and the associated absence of CpG hypermutability (Raddatz et al. 2013). In combining multiple detection strategies with stringent filtering steps, including manual inspection of calls we are confident that our final call set comprises true somatic mutations in ISCs.

We first considered all genome-wide mutations, and found a median of 44 and 123 SNVs and indels per sample, respectively (Fig. S4b, c), approximately 1.4 somatic mutations per megabase. Strikingly, this mutation prevalence of 1.4 per Mb is broadly similar to those typically found in several human cancers such as ovarian (1.85 per Mb) and breast (1.29 per Mb) (Greenman et al. 2007; Alexandrov et al. 2013a; Angus et al. 2019). Considering that tumours were dissected from six-week-old files, this suggests an overall high mutation rate in flies relative to human cancer genomes. We find no evidence of kataegis - hypermutation in localised genomic regions - which is sometimes found in cancer genomes (Alexandrov et al. 2013a). Next, to focus on mutations in protein-coding regions, we combined SNV and indel calls and annotated mutations with their functional impact (Martincorena et al. 2017), and found a median of 12 protein-coding mutations per sample (Fig. 4d; Supplementary Table 3). Of particular interest, our analysis did not uncover any protein-coding mutations in *Notch*, or in components of the Notch signalling pathway.

Next, to gain insight into the selective dynamics operating within tumours, and to identify other potential drivers of tumour formation, we calculated the ratio of non-synonymous substitutions to synonymous substitutions (dN/dS). While we see several genes with multiple protein-coding mutations in different samples, we do not find statistical support for any of these being potential drivers. Interestingly, our dN/dS analysis revealed a global ratio of 0.34, indicative of negative selection, which deviates substantially from values that are typical of both human cancer genomes and somatic tissues (Martincorena et al. 2017). While we discuss interpretations of this observed difference later, such comparisons between phyla are important to assess the universality of selective dynamics operating in somatic tissues.

Human cancer genomes frequently exhibit patterns of mutation - mutational signatures - that are often associated with exposure to distinct underlying mutational processes, which have been extensively catalogued in large-scale analyses (Alexandrov et al. 2013a, 2020, 2013b; Nik-Zainal et al. 2012), and categorised into signatures in the COSMIC database (https://cancer.sanger.ac.uk/cosmic/signatures_v2). In order to investigate whether we could discern any underlying mutational processes influencing the spectrum of mutations we observe, we first extended our SNV analysis to consider mutations within a trinucleotide (the immediate 5’ and 3’ bases) sequence context. Combining data from all samples, we observe that T>C and C>T transitions were marginally more frequent than C>A, C>G, T>A, T>G transversions, however we do not find any context-specific enrichment of distinct 5’ and 3’ bases associated with transitions (Fig. S4d). Next, to examine mutational patterns operative within individual samples, we then calculated the per-sample cosine similarity between mutational profiles and COSMIC signatures (Blokzijl et al. 2018). While we did not find any one signature contributing to large numbers of mutations across samples, we identified several signatures that contributed heavily in several samples (Fig. S4e). Signature 3 was observed to contribute strongly in five samples (P45a, P47, P63, D1 and D7), which has previously been found to be associated with BRCA1 deficiency and failure of DSB-repair by homologous recombination in human breast cancer genomes (Nik-Zainal et al. 2016). Interestingly, we also see a relatively strong contribution of signature 16 in five different samples (P9, P17, P37, P53 and P55), which is a liver-cancer specific signature, highly associated with alcohol consumption (Letouzé et al. 2017). Several other signatures (5, 8) were detected in other samples for which the underlying mutational processes remain unknown. Importantly, however, since these signatures are found in flies as well as humans, it is likely the biological processes that generate such signatures are also operative in the fly gut.

### 5. Genome-wide distribution of mutations

In human cancer genomes, several features contribute to the non-random distribution of structural variant breakpoints, SNVs and indels, including local base composition, chromatin structure, and gene expression (Schuster-Böckler and Lehner 2012). To determine the extent to which mutations are enriched or depleted in a given genome feature, we compared the number of mutations observed in the feature to the number of mutations expected in the feature by chance considering its total length. As one particular sample (P7; Fig. 4c, Fig. S4b, c) contributed heavily to the total mutational burden across all samples, we excluded this sample from all subsequent aggregate analyses to avoid sample-specific bias.

First, we concentrated on the mutations in both coding (CDS) and non-coding (UTR, introns) gene features. Overall, we found all classes of mutation to be weakly depleted in genic regions of the genome (Fig. 5a), consistent with the observation that euchromatic regions are depleted for mutations (Pleasance et al. 2010; Woo and Li 2012; Schuster-Böckler and Lehner 2012). However, we find CDS to be strongly depleted for both SNVs and indels, suggesting that such regions may be maintained under negative selection. In order to investigate this further, we annotated our data to include expression levels using recently published ISC-specific RNA-Seq data (Dutta et al. 2015). Interestingly, we found that the coding sequences of ISC-expressed genes were more strongly depleted for indels, but not SNVs, than in non-expressed genes (Fig. S5a). We also observed that both 3’ and 5’ UTRs were approximately 3 fold more strongly enriched for mutations in expressed versus non-expressed genes. Considering mutations in UTRs have been traditionally overlooked by studies focussing on protein-coding regions of the genome, and that a subset of highly expressed oncogenes are frequently mutated at their 3’UTR in human cancer genomes (Supek et al. 2014), this finding highlights that mutations in UTRs may be an important, but under-investigated, class of mutation.

**Fig. 5.**
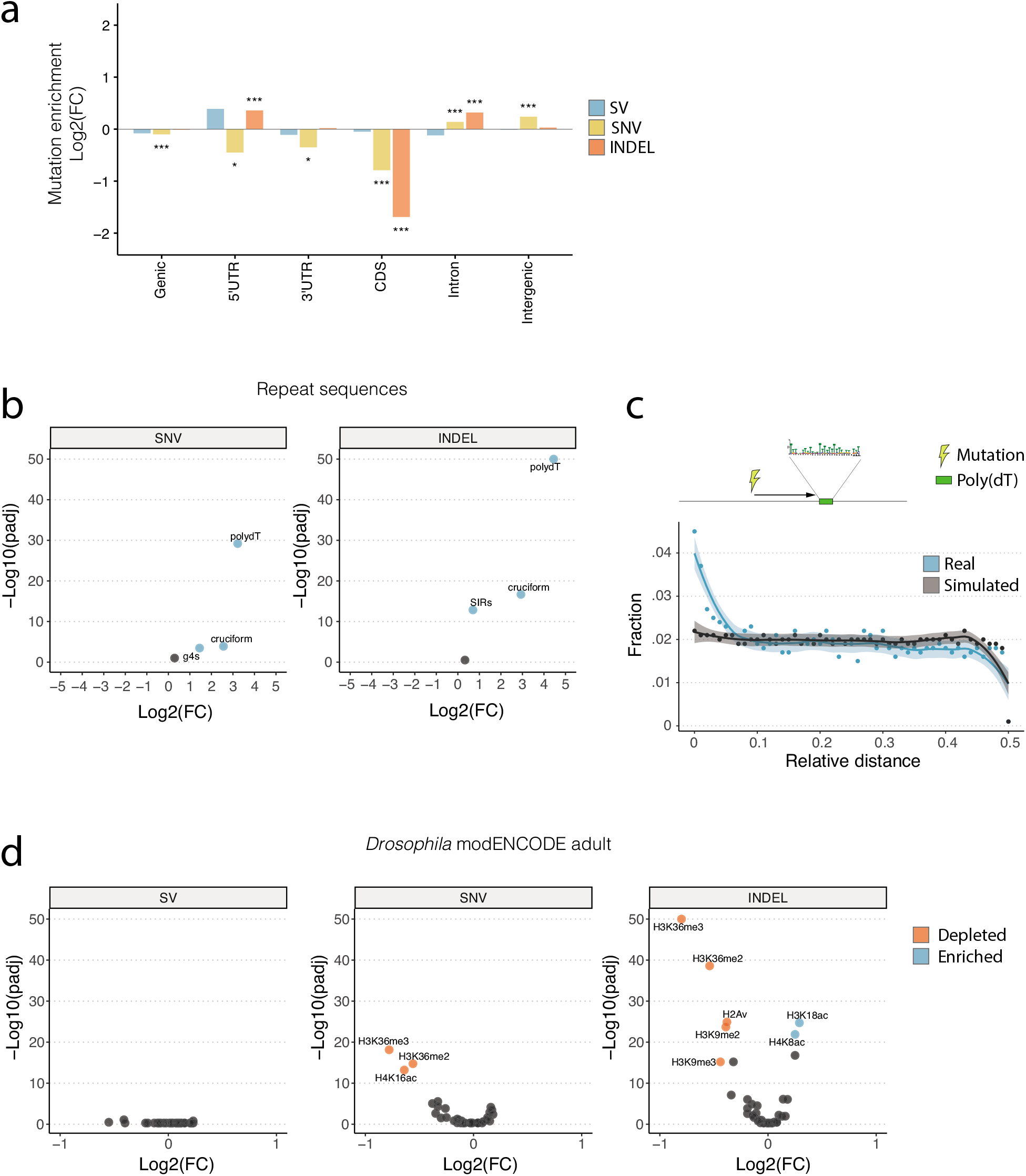
Distribution of somatic mutations in genome features. (**a**) Point mutations (SNVs and indels) were both strongly depleted in CDS regions. (**b**, **d**) Volcano plots showing enrichment or depletion of mutations in repeat regions (**b**) and chromatin features from the *Drosophila* modENCODE dataset (**d**). Highlighted features represent those that with an E-score (-Log10(p)*Log2(FC); Methods) > 5. (**c**) The distribution of relative distances between combined somatic mutations (breakpoints, SNVs and indels; shown in blue) and the closest instance of poly(dA:dT) tracts in the genome. Simulated data are shown for comparison in grey. The y axis of **b** and **d** are restricted to a maximum - Log10(padj) value of 50. Asterisks denote significance: *** p < 0.001; ** p < 0.01; * p < 0.5. All p values shown have been generated from a two-sided binomial test, and adjusted for multiple comparisons using a Benjamini-Hochberg adjustment.

Repeat sequences have previously been associated with increased mutability, and have been shown to be enriched around structural variant breakpoints (Lu et al. 2015) and for point mutations (Zou et al. 2017) in human cancers. Consistent with such reports, we found a strong enrichment for point mutations in inverted repeats, which was particularly notable in cruciform DNA (Fig. 5b), suggesting that similar mutational dynamics are operative in the fly genome.

Interestingly, we also found an extreme genome-wide enrichment of point mutations in the poly(dA:dT) tracts (Fig. 5b). In comparing the relative distance of mutations to poly(dA:T) tracts (Methods), we find that not only are mutations enriched in poly(dA:dT) tracts, but that they are also found closer to such repeats more frequently than in randomly distributed data (Fig. 5c), suggesting that such repeats are both inherently mutable and play a role in determining the mutation rate of flanking DNA.

Considering that chromatin organisation has also been shown to influence mutation in cancer genomes (Schuster-Böckler and Lehner 2012), we next investigated the distribution of mutations in the publicly available *Drosophila* modENCODE adult fly datasets, including chromatin landscape and transcription factor binding sites (modENCODE Consortium et al. 2010). We found a strong depletion for both SNVs and indels in chromatin regions enriched for several marks including H3K36me2/3 and H3K9me2/3, which is associated with transcriptional repression (Fig. 5d). In contrast, SNVs in cancer genomes are enriched in H3K9me2/3 (Schuster-Böckler and Lehner 2012), suggesting that these marks may influence mutational processes differently in *Drosophila*. Conversely, we found an enrichment of indels in marks associated with transcriptional activation (H4K8ac, H3K18ac) and several transcription factor binding sites, all of which belonged to either the C2H2 family of zinc-finger proteins (br, Trl, Cf2, odd, hb) or HMG proteins (pan, D; Fig. S5b). Interestingly binding sites of CTCF, a well-known member of the C2H2 family, have been found to be highly enriched for point mutations in cancer genomes (Katainen et al. 2015). While we do not find an enrichment of mutations in CTCF sites, it is possible that other C2H2 family members are involved in similar processes, and further work is required to establish the significance of this finding.

Finally, in order to establish whether a similar distribution of mutations was observable in ISC-specific chromatin profiles, we repeated enrichment analyses using our recently published DamID profiles of chromatin binding factors in ISCs (Gervais et al. 2019). In agreement with the results found in the *Drosophila* modENCODE datasets, mutations were found to be depleted in regions associated with silent chromatin, marked by Heterochromatin Protein 1 (HP1), and enriched in regions bound by Trithorax-related (Trr), RNA polymerase II (PolII) and Kismet (Fig. S5c), all of which have been previously shown to map transcriptionally active chromatin (Gervais et al. 2019).

Thus, somatic mutations in ISCs are distributed non-randomly across the *Drosophila* genome, and are found associated with features that influence mutation distribution in cancer genomes, as well as with features with no known associations. Taken together, these findings demonstrate the necessity of exploring mutation distribution across whole genomes, and the value of performing such analyses in *Drosophila*.

### 6. Mutational timing and tumour evolution

Each tumour genome bears the cumulative damage acquired over its evolutionary history, which can be partially reconstructed using whole-genome sequencing. Mutations arising early in adult life will be propagated throughout the ISC lineage whereas those arising after tumour formation will be subclonal to the driving mutation and present in a smaller fraction of cells (Fig. 6a, b). Considering the time frame of our experiment, we estimate that the mutations driving *Notch* inactivation arise at around three weeks post eclosion, and that tumours then develop for another three weeks before dissection (Siudeja et al. 2015). To reconstruct the evolutionary history of each tumour, we treated the variant allele frequency (VAF) of each variant as a proxy for mutational time. To normalise between mutations on sex chromosomes and autosomes, we multiplied the VAF on autosomes by a factor of two. As the origin of each tumour we have sequenced can be explained by a mutational event affecting the Notch signalling pathway, we approximated mutational timing relative to *Notch*-inactivating events.

**Fig. 6.**
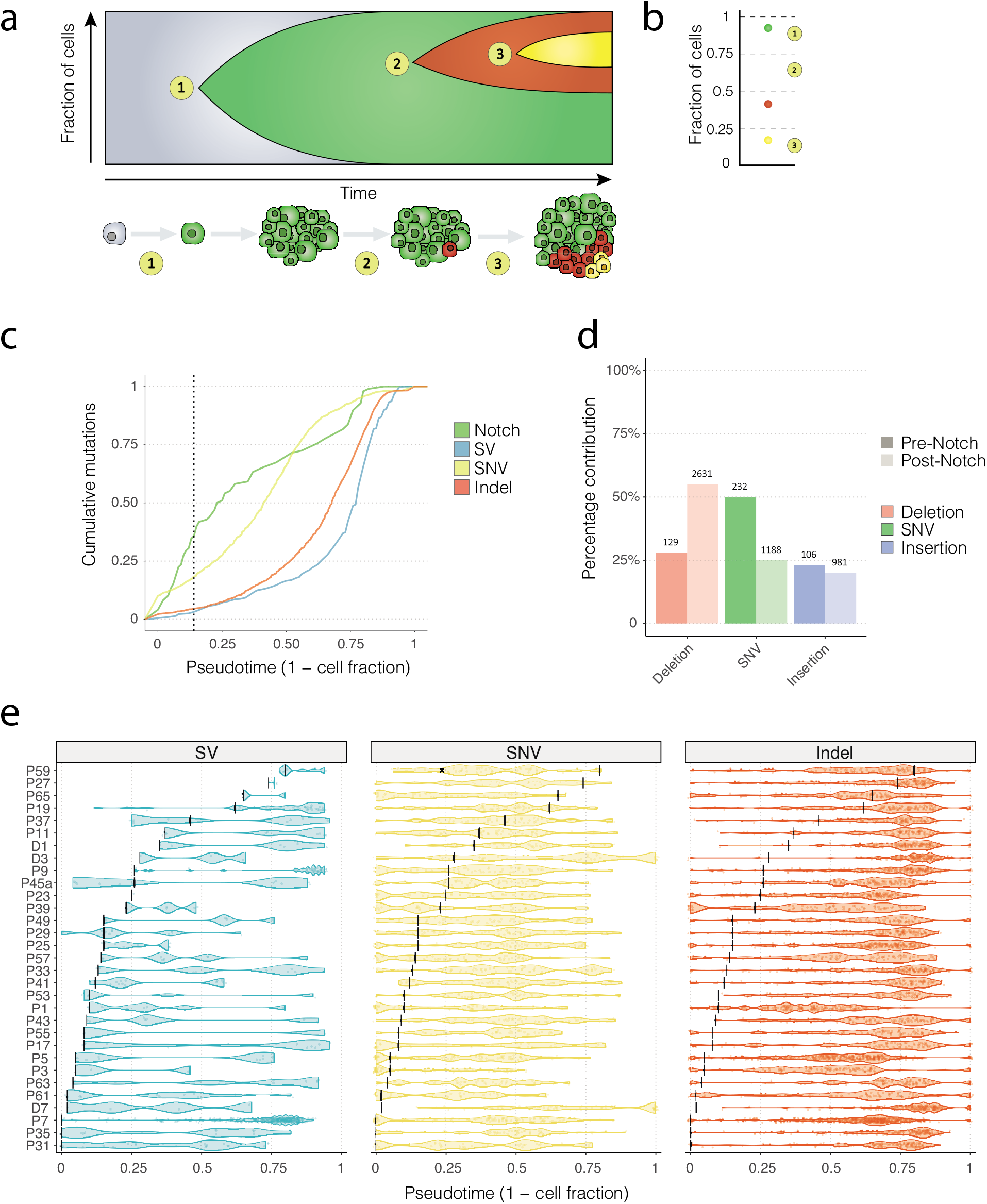
The evolution of somatic mutational in ISC genomes. (**a**) A schematic illustrating the accumulation of mutations in a stem cell and neoplastic clone over time. At time point 1, an ISC acquires a somatic mutation that inactivates *Notch*, driving hyperproliferation of *Notch-* cells (in green). At time points 2 and 3, subsequent mutations are acquired (shown in red and yellow) that are present in smaller numbers of cells. (**b**) A modified VAF (Methods) can be used to estimate the fraction of cells carrying each mutation that we use as a proxy for time (pseudotime). (**c**) The cumulative distribution of mutations (aggregated over all samples) over pseudotime show that mutations in *Notch* occur before other classes of mutation. SNVs arose prior to indels and additional SVs. The median pseudotime value for *Notch*-inactivating events is shown as a dotted vertical line. (**d**) For each sample, VAF values of *Notch*-inactivating mutations were used to divide point mutations observed genome-wide as occurring *pre-Notch* (darker shaded bars) and post-*Notch* (lighter shaded bars). Numbers on top of each bar show the number of mutations observed in each category. (**e**) Per-sample estimates of tumour evolution. *Notch*-inactivating events for each sample are shown as vertical black bars. Each dot represents a single mutation, and violin plots ease the visualisation of mutation distribution over pseudotime.

Consistent with the notion that *Notch* mutations occur early in tumour evolution, we find the majority of all mutations across samples to be subclonal to the mutation in *Notch* (84%; 4664/5546; Fig. 6c). This would also suggest that cells in the tumour experience a higher mutational burden than those in pre-tumour ISCs. However, we found 8.8% of point mutations to occur prior to the mutation in *Notch* (467/5267), that presumably took place during development or in young adult ISCs. In comparing frequencies of mutation types between pre- and post-*Notch* mutations, we find differences in the frequencies of small deletions and SNVs, but not small insertions (Fig. 6d). Interestingly, we find that small deletions comprise 29% of point mutations prior to *Notch*-inactivation, and 57% of those occurring after. This observation is even more pronounced in protein-coding mutations, where we do not detect any frame-shift-inducing small deletions *pre-Notch* (Fig. S6b).

Interestingly, in many samples the majority of indels arose late in tumour evolution, suggestive of a sudden burst of mutation (Fig. 6e). Such rapid accumulation of mutations is frequently observed in human cancer genomes after the loss of a DNA-damage repair component. To assess whether this might explain the observed VAF distribution, we searched for mutations affecting genes in DNA-damage repair pathways. While we found several mutations in genes that could potentially explain increased rates of mutation, only one of them (R59; Fig. 6e) - a missense mutation in the *Drosophila Rad51* homolog - had coding consequences. This suggests that mutation of DNA-damage repair components is not responsible for the accumulation of mutations in late tumour evolution *per se*. It is however possible that such late-occurring mutations are a consequence of other processes, such as age-related changes in DNA repair efficiency or mutations in the non-coding genome, such as regulatory elements of DNA repair genes.

Together, our findings indicate that in this model system of spontaneous neoplasia formation in wild-type flies, the major driver events are loss of *Notch* activity through deletions and complex rearrangements. Subsequent genome diversity then arises via the accumulation of SNVs, indels, and additional structural variants. Our data demonstrate a surprisingly rapid accumulation of mutations over a short evolutionary time span during the adult life of *Drosophila melanogaster*.

## Discussion

We have applied whole-genome sequencing to interrogate how spontaneous tumours arise from stem cells in the *Drosophila* intestine. Our in-depth analysis of the causes of driver inactivation and subsequent tumour evolution, provide insight into spontaneous somatic mutation over short time-scales.

### Evidence for a replicative mechanism underlying *Notch* inactivation

By analysing the mutations affecting *Notch*, we have shown compelling evidence for the involvement of a replicative mechanism in driving tumour formation. Firstly, 23% of the *Notch*inactivating events were classified as complex rearrangements, involving multiple connected breakpoint junctions, many of which harboured inserted sequences. Both of these observations are best explained by the involvement of a replicative mechanism, such as MMBIR. Secondly, sequences flanking the breakpoints of *Notch*-inactivating structural variants were found to be highly enriched for poly(dA:dT) repeats that have previously been implicated as preferential loci for replication fork stalling and collapse after induction of replication stress (Tubbs et al. 2018). Interestingly, there was a strong enrichment for point mutations in both poly(dA:dT) sequences and inverted repeats, and mutations were found to occur closer to repeat sequences than expected by chance. One explanation for this association is that repeat sequences are prone to adopting non-B-form DNA structures, which can stall replication forks, causing replication fork collapse and exposing highly mutable singlestranded DNA (Lu et al. 2015; Kurahashi et al. 2004; Tubbs et al. 2018).

### Genome-wide distribution and mutational burden in ISCs

By exploiting the clonal nature of the tumour, we were also able to characterise somatic mutations genome-wide. Overall, highly subclonal translocations were the most frequent type of structural variant detected across the genome, although these largely originated from one sample (P7). It is important to note that the read-depth-based approach to detecting CNVs are inherently less sensitive than read-mapping-based approaches, and as such, we expect to detect copy-number-neutral variants (such as translocations and inversions) at lower VAFs than CNVs. In addition, considering that our pipeline integrates both read depth and frequency changes of heterozygous SNPs to filter CNVs, we have less information with which to discern false positive events for copy-number-neutral variants. However, we found both the *Notch* variants and those detected genome-wide to be highly deficient for tandem duplications, a class of structural variant commonly found in both cancer (Li et al. 2020), and normal somatic genomes (Lee-Six et al. 2019; Moore et al. 2020). Unlike the structural variants in *Notch*, this difference observed in genome-wide variants cannot be explained by an influence of our experimental setup (which selects for *Notch*-inactivating events), suggesting the intriguing possibility of a mechanistic difference for generating tandem duplications in *Drosophila*.

### Tumour evolution

Using a tumour-purity-adjusted VAF, we attempted to reconstruct the evolutionary history of somatic mutation in ISC genomes. While similar approaches have been used recently and applied to human cancer data (Gerstung et al. 2020), it is important to note the potential limitations of this strategy. First of all, explicitly using VAF as a proxy for mutational timing assumes a linear propagation of mutations within clones, which will almost certainly fail to capture the dynamics of clone contraction/expansion operating within a tumour. Here, a mutation that occurs very early in tumour development may be selected against in the tumour, and be present in few cells of the dissected tissue. The approach we take is therefore unable to distinguish early-occurring, negatively selected, mutations from those that are late-occurring and positively selected. In addition, while it has little bearing on our interpretation, it is unable to distinguish mutations in separate cell populations from those occurring within the same clone. Nonetheless, by estimating the timing of mutations in this fashion, we were able to establish the *Notch* events as founding mutations, and divide other somatic mutations into those occurring pre- and post-*Notch* inactivation. We find that 8.8% of point mutations (and 5.4% of those in protein-coding domains) occur before *Notch* inactivation, in normal cells. In comparing the relative proportions of mutation types in pre- and post-*Notch* groups we observed that pre-*Notch* mutations were relatively depleted for short deletions, and enriched for SNVs. This relationship was also present in point mutations in protein coding regions, with no frame-shift inducing short deletions observed pre-*Notch*. Whether this is due to negative selection on more deleterious mutations or a mechanistic difference between early and late tumour development remains to be seen. However, we have shown that the majority of mutations occur post-*Notch* inactivation, consistent with an accelerated rate of mutation in tumours. This could be due to an increased number of cell divisions following neoplasia formation, or changes in the mutation rate arising during ageing or tumour development.

### A novel insight into selective dynamics in somatic tissues

On extending our analysis to include point mutations across the whole genome, we found a relatively high frequency of mutation (1.4 per Mb), a mutation prevalence comparable to several human cancers (Greenman et al. 2007; Alexandrov et al. 2013a; Angus et al. 2019). Considering that we dissect tumours from flies at around six weeks post-eclosion, and that the tumour itself has only been developing for around three weeks, this would imply that in a matter of weeks, *Drosophila* ISCs reach a mutational burden equivalent to several decades worth of mutation in human cancers. On screening for mutations in genes that could drive high mutation rates (such as DNA-damage repair components) we only found one sample harbouring a protein-coding mutation in the *Drosophila* Rad51 homolog, suggesting that the mutation burden observed across samples does not result from genetic impairment in the ability to repair damage to DNA. In agreement with this, our mutational signature analysis did not identify signatures associated with defective DNA mismatch repair (COSMIC v2 signatures 6, 15, 20, and 26).

Comparative genomic analyses of closely related species reveal that species-level evolution is characterised by negative selection, where the majority of germline non-synonymous mutations are selected against during species evolution (Ostrow et al. 2014). In mammals however, both somatic tissues and cancer genomes are characterised by positive selection, where the ratio of non-synonymous to synonymous mutations (dN/dS) is greater than 1 (Martincorena et al. 2017). In contrast to these observations, we find a dN/dS ratio of 0.34, suggesting that ISC genomes are maintained under negative selection. One explanation for this would be if the somatic point mutations identified included germline variants, which would drive down the somatic dN/dS ratio. However, considering that we include all normal samples in our panel of normals, and stringently remove variants found in any normal sample, this explanation seems unlikely. Another possibility is that the relative paucity of paralogous genes in the *Drosophila* genome, compared to vertebrate genomes, means that there is stronger selective pressure to maintain gene function in somatic cells. While it would seem likely that this would be somewhat negated by having two alleles on autosomes, it is an interesting observation that challenges the current paradigm of selection in somatic cells. Finally, considering the very short time period with which we detect mutations - approximately three weeks pre- and post-*Notch* inactivation (Siudeja et al. 2015) - it is possible that the normal selective dynamics operative in somatic tissues do not have enough time to exert a detectable effect on the cells carrying protein-coding mutations. If this were the case, it would suggest that our assay affords us rare insight into processes operating in the very early stages of cancer, that are absent from human cancer studies.

## Methods

### Sequencing of Drosophila neoplasia and controls

A detailed methodology can be found in (Siudeja et al. 2020). In brief, for selection of neoplastic tissue, clusters of EE cells or ISC cells were selected by expression of Prospero-Gal4- (*Pros>2XGFP*) or Dl-Gal4- (*Dl>Gal4*) driven UAS-GFP (*Pros>2XGFP*) adult flies were obtained by crossing *w-;;Pros^Volia1^GAL4/TM6BTbSb* females (gift from J. de Navascués) with *w-;UAS-2XGFP*; males (Bloomington). *Dl>nlsGFP* flies were obtained by crossing *w-;;DlGal4/TM6TbHu* (gift from S. Hou) females with *w-;;UAS-nlsGFP* males (Bloomington). 6-7-weeks-old *Pros>2XGFP* or *Dl>nlsGFP* males were used to visually identify midguts containing neoplasia based on clonal accumulation of GFP positive cells. To isolate neoplasia, the midgut region containing an estimated 40%–80% neoplastic cells was manually dissected together with the head as a direct comparison. Genomic DNA for short-read Illumina sequencing was isolated with the QIAamp DNA MicroKit (Qiagen) according to the manufacturer’s protocol dedicated to processing laser-microdissected tissues. DNA quantity was measured with Qubit dsDNA High Sensitivity Assay Kit. For three tumour normal pairs, data were reanalysed from our previous work (Siudeja et al. 2015).

### DNA sequencing

Genomic DNA libraries were prepared with the Nextera XT protocol (Illumina). Whole-genome 2X100 bps or 2X150 bps paired-end sequencing was performed on HiSeq2500 or Novaseq (Illumina) on a total of 34 samples and their respective head controls (Supplementary table 4). Two samples (P13, P21) were excluded from further analysis due to low coverage.

### Point mutation calling and filtering

Reads were aligned to the *Drosophila* genome release 6.12 using bwa mem v0.7.15, and duplicate reads were marked using Picard MarkDuplicates v2.7.1 (http://broadinstitute.github.io/picard/). samclip (https://github.com/tseemann/samclip) was used to remove clipped reads. A PON was constructed by running Mutect2 (v4.1.2; (Cibulskis et al. 2013)) on all normal samples, and combining them using CreateSomaticPanelOfNormals from the GATK suite of tools (v4.1.2; (McKenna et al. 2010)). We then called somatic point mutations using several different approaches. For each sample we ran Mutect2 (v4.1.2), Varscan2 (v2.4; (Koboldt et al. 2012)), Strelka (v2.9.10; (Kim et al. 2018)), SomaticSniper (v1.0.5.0; (Larson et al. 2012)) in tumour/normal mode and merged calls per-sample using SomaticSeq (v3.3.0; (Fang et al. 2015)). Freebayes (v1.2.0-dirty; (Garrison and Marth 2012)) was then run in somatic mode, and we merged output from SomaticSeq with Freebayes calls to create a unified per-sample call set. These calls were then filtered against the PON to remove germline variants and select for variants called at regions with read depth >= 20 in both the tumour and normal sample (Supplementary Methods).

### Point mutation annotation

Initially, we annotated point mutations with gene information using SnpEff v4.3 (Cingolani et al. 2012), and used ISC-specific RNA-Seq data (Dutta et al. 2015) to add expression levels to point mutations in genes. We then used the R package dNdScv v0.1 (Martincorena et al. 2017) to annotate protein-coding mutations for their functional impact.

### Mutational Signatures analysis

We used MutationalPatterns v1.8 (Blokzijl et al. 2018) to detect relative contributions made by mutational signatures in the COSMIC database (https://cancer.sanger.ac.uk/cosmic/signatures_v2). Filtered vcf files were used to generate a matrix of mutational counts at trinucleotide positions and the fit_to_signatures function was used to assign the optimal linear combination of mutational signatures that most closely explained the per-sample mutational spectrum. Contributions were plotted using the plot_contribution_heatmap function.

### Read tagging

We have recently developed a bioinformatic tool named readtagger to tag reads that map to multiple genomes ((Siudeja et al. 2020)). In cases where reads map to a primary genome with less affinity than to a secondary genome, reads are tagged as originating from the secondary source, and can then be filtered or extracted from the alignment to the primary genome. Using this approach, we first mapped reads to *Drosophila* reference TE sequences using bwa mem v0.7.15 to annotate non-reference TE sequences in the genomes of our fly stocks. Importantly, this step enables the easy extraction and annotation of clusters of transposable element-mapped reads in downstream analysis. Next, in order to filter out reads that likely originated from contaminating species of the *Drosophila* microbiome, we mapped reads to several known species found in the *Drosophila* gut *(Acetobacter pasteurianus* (NC_013209.1), *Escherichia coli* (NC_000913.3), Innubila nudivirus (NC_040699.1), *Komagataeibacter nataicola* (NZ_CP019875.1), *Lactobacillus brevis* (NC_008497.1), *Lactobacillus plantarum* (NC_004567.2), *Plasmodium yoelii* (LR129838.1), *Saccharomyces* (NC_001133), Tomelloso virus (KY457233.1), *Wolbachia pipientis* (NC_002978.6)). Finally reads were mapped to *Drosophila melanogaster* genome release 6.12, and readtagger v0.4.11 was used to tag reads. Duplicate reads were marked using Picard MarkDuplicates v2.7.1 (http://broadinstitute.github.io/picard/).

### Structural variant calling and filtering

First, CNVs were called using two different approaches CNV-Seq (Xie and Tammi 2009) and Control-FREEC v11.0 (Boeva et al. 2012). Next, we used three different read-based approaches to precisely identify breakpoints: novoBreak v1.1 (Chong et al. 2017), LUMPY v0.2.13 (Layer et al. 2014) and DELLY v0.7.8 (Rausch et al. 2012). We created a PON by genotyping all normal samples using SVTyper v0.0.4 (Chiang et al. 2015) and used these to remove germline calls. We then used the tool svParse v0.3.1 developed within the svParser suite of tools (https://github.com/bardin-lab/svParser) to filter variants, selecting for those in the mappable genome, supported by at least 3 reads, with a read depth of at least 10 in both the tumour and normal, and a ratio of read depth to supporting reads > 0.05. The output of CNV-Seq was used to annotate all variants with a Log2(FC) tumour/normal read depth ratio and we combined all calls per sample, clustering variants called by multiple approaches into the same mutational event. We then annotated breakpoints in genes for expression levels using ISC-specific RNA-Seq data (Dutta et al. 2015). In order to assign a putative underlying mechanism to each structural variant, breakpoints that were supported by multiple split-reads were annotated with potential microhomology sequences using SplitVision (Nazaryan-Petersen et al. 2018). Next, we attempted to re-align any sequences inserted at breakpoint junctions to the sequence flanking each breakpoint to determine whether they were locally templated. We then categorised structural variants using criteria largely adapted from (Yang et al. 2013; Kidd et al. 2010) (Fig. S1b; Supplementary Tables 1 and 2).

### Breakpoint recalibration

Per-sample call sets were re-calibrated and further filtered using svSupport (https://github.com/bardin-lab/svSupport). In this step we removed variants supported by unannotated duplicate reads and standardised annotation of variants according to read mapping signatures. In order to unify variants identified by multiple approaches we performed a local search for split-read support (Supplementary Methods), and adjusted breakpoints accordingly. We then extracted reads that supported and directly opposed each variant, and used these read counts to calculate variant allele frequency (VAF) by dividing the number of supporting reads by the number of supporting reads + number of opposing reads. Tumour purity values obtained from Control-FREEC were then used to calculate a tumour-purity adjusted value by adjusting the number of reads expected to oppose each variant, given a per-sample purity value. For example, a variant supported by 75 reads, and opposed by 25 reads with a sample tumour purity value of 0.75 would have an initial VAF of 0.75 (75/(75 + 25)), and an adjusted VAF of 1 (supporting/(supporting + (opposing - (1 - purity) * total reads)); 75/(75 + (25 - (1 - 0.75) * (75 + 25))). Importantly, this step enables us to better estimate the timing of mutations.

### Breakpoint clustering

To record complex rearrangements as a distinct class of structural variant, we clustered breakpoints within +/- 5 kb and merged them into a single event using svStitch (https://github.com/bardin-lab/svParser/blob/master/script/svStitch.py). In cases where clustered variants all belonged to the same class of CN event (deletion or duplication) individual variants were collapsed into a single event and the variant type was not modified. In cases where multiple classes of structural variant class were clustered, or all classes belonged to the same class but were not CN events, we re-annotated each variant type in the cluster as “complex”, and collapsed variants into a single mutational event.

### Annotating CNVs with SNP allele frequency

To further improve our ability to discern false-positive CNV events in our structural variant data, we used germline SNP allele frequencies to detect shifts in copy number over CNV regions. For each sample, we used calls made by Freebayes to compare the allele frequency of germline SNPs in regions marked as being CNV without split read support. For cases where the average allele frequency shift over the CNV was below 10% and the number of informative sites surveyed was fewer than 5 we marked CNVs as false positives, and excluded them from further analysis.

### Manual inspection of *Notch* variants

For each sample in *Notch*, we manually inspected breakpoints using IGV. In several samples, we manually added CNVs that were below the detection threshold of our pipeline, and in cases where breakpoints were unresolved, we adjusted these in line with split-read evidence where possible. In one sample (P59) we manually reclassified a complex event as two distinct inversions, as we could not find any read evidence linking the breakpoints. We then used IGV to identify TE-tagged reads at breakpoints, and manually characterised the signature of TE-tagged reads for each variant in *Notch*. Here, we classified TE presence at breakpoints as either somatic or germline, and recorded the TE family best supported by clusters of TE-tagged reads.

### Identification of Tomelloso virus at breakpoint junctions and in sequencing data

In one sample (P31) we detected reads at a breakpoint junction in *Notch* whose mates were not mapped to the *Drosophila* genome. In order to detect the source of these unmapped reads, we extracted unmapped reads, and assembled them into contigs using CAP3 (Huang and Madan 1999). We then used the assembled contigs to query the nr database using BLASTn (Altschul et al. 1990), and identified dsDNA nudivirus Tomelloso (Palmer et al. 2018) as the source. We then extended this search genome-wide, by identifying somatic clusters of reads with unmapped mates and assembling their mates as described above, but found no other instances of viral integration in other samples. In order to assess the extent to which Tomelloso was present in our sequencing data, we combined the *Drosophila* and Tomelloso genomes tagged reads that likely originated from Tomelloso using readtagger v0.4.11. We then analysed the percentage of paired-end reads that mapped with high confidence to the Tomelloso genome in all samples, and performed a Pearson correlation to determine the relationship between mutation per-sample mutation count and viral load. These analyses excluded a relationship between viral load and mutation frequency.

### Genome feature discovery

To identify regions of the *Drosophila* genome release 6.12 susceptible to forming non-B-form DNA structures we first scanned the genome for inverted repeat sequences following the approach outlined in (Zou et al. 2017), as well as for G-quadruplexes using the R package G4Hunter (Bedrat et al. 2016). To identify short sequence motifs in *Notch* breakpoint regions, we used MEME v5.0.1 (Bailey and Elkan 1994), using sequences extracted +/- 500 bps from each breakpoint in *Notch* as input. We then used FIMO v5.0.1 (Grant et al. 2011) to search for and annotate recovered motifs genome-wide.

### Association of mutations with genomic regions

To detect the enrichment or depletion of genomic regions for mutations we counted the number of mutations in a given region, and compared this to the expected number considering the region’s size. The association was tested by performing a two-sided binomial test, adjusting for multiple comparisons using Benjamini-Hochberg adjustment. We require that the number of observed hits + expected hits is greater than 10 for plotting. To assess whether breakpoints in *Notch* were enriched for poly(dA:dT) sequences, the sequence +/- 500 bps around each breakpoint in *Notch* was extracted and permutation tests were performed using regioneR v1.14 (Gel et al. 2016) on the overlap between repeats and these breakpoint-flanking regions. In order to compare observed counts between real and shuffled data, we restricted permutations to within the genomic locus X:2700000-3400000, and performed 10,000 permutations. To test whether mutations were found closer to poly(dA:dT) tracts than expected by chance, all classes of mutation (structural variant breakpoints, SNV and indels) were combined. We then simulated mutations across the mappable genome with distribution across chromosomes equal to that observed in our combined mutation data to act as a comparison. Finally, we calculated the relative distances of both mutations and simulated data to the closest repeat sequence using bedtools reldist v2.28 (Favorov et al. 2012).

## Declarations

### Ethics approval and consent to participate

Not applicable

### Consent for publication

Not applicable

### Authors’ contributions

NR developed software, analysed and interpreted the data, and wrote the manuscript. KS prepared samples, and helped interpret data. MvdB developed readtagger and helped interpret data. BB helped prepare samples and helped interpret data. AJB, conceived and supervised the study, interpreted the data, and wrote the manuscript.

## Acknowledgements

We thank P-A. Defossez, N. Servant, J. Waterfall, and members of the Bardin lab for helpful discussions, critiques, and comments on the manuscript and J. de Navascués, S. Hou, and the Bloomington stock centre for Drosophila stocks.

## Availability of data and materials

The dataset supporting the conclusions of this article is available in the NCBI Sequence Read Archive (SRA) database under BioProject PRJNA641572. The three tumour normal pairs previously reported (Siudeja et al. 2015) are available under the ArrayExpress accession number E-MTAB-3917.

## References

Alexandrov LB, Kim J, Haradhvala NJ, Huang MN, Tian Ng AW, Wu Y, Boot A, Covington KR, Gordenin DA, Bergstrom EN, et al. 2020. The repertoire of mutational signatures in human cancer. Nature 578: 94–101.

Alexandrov LB, Nik-Zainal S, Wedge DC, Aparicio SAJR, Behjati S, Biankin AV, Bignell GR, Bolli N, Borg A, Børresen-Dale A-L, et al. 2013a. Signatures of mutational processes in human cancer. Nature 500: 415–421.

Alexandrov LB, Nik-Zainal S, Wedge DC, Campbell PJ, Stratton MR. 2013b. Deciphering signatures of mutational processes operative in human cancer. Cell Rep 3: 246–259.

Altschul SF, Gish W, Miller W, Myers EW, Lipman DJ. 1990. Basic local alignment search tool. J Mol Biol 215: 403–410.

Al Zouabi L, Bardin AJ. 2020. Stem Cell DNA Damage and Genome Mutation in the Context of Aging and Cancer Initiation. Cold Spring Harb Perspect Biol. http://dx.doi.org/10.1101/cshperspect.a036210.

Angus L, Smid M, Wilting SM, van Riet J, Van Hoeck A, Nguyen L, Nik-Zainal S, Steenbruggen TG, Tjan-Heijnen VCG, Labots M, et al. 2019. The genomic landscape of metastatic breast cancer highlights changes in mutation and signature frequencies. Nat Genet 51: 1450–1458.

Bailey MH, Tokheim C, Porta-Pardo E, Sengupta S, Bertrand D, Weerasinghe A, Colaprico A, Wendl MC, Kim J, Reardon B, et al. 2018. Comprehensive Characterization of Cancer Driver Genes and Mutations. Cell 173: 371–385.e18.

Bailey TL, Elkan C. 1994. Fitting a mixture model by expectation maximization to discover motifs in biopolymers. Proc Int Conf Intell Syst Mol Biol 2: 28–36.

Bainbridge MN, Wang M, Wu Y, Newsham I, Muzny DM, Jefferies JL, Albert TJ, Burgess DL, Gibbs RA. 2011. Targeted enrichment beyond the consensus coding DNA sequence exome reveals exons with higher variant densities. Genome Biol 12: R68.

Bedrat A, Lacroix L, Mergny J-L. 2016. Re-evaluation of G-quadruplex propensity with G4Hunter. Nucleic Acids Res 44: 1746–1759.

Blokzijl F, Janssen R, van Boxtel R, Cuppen E. 2018. MutationalPatterns: comprehensive genome-wide analysis of mutational processes. Genome Med 10: 33.

Boeva V, Popova T, Bleakley K, Chiche P, Cappo J, Schleiermacher G, Janoueix-Lerosey I, Delattre O, Barillot E. 2012. Control-FREEC: a tool for assessing copy number and allelic content using next-generation sequencing data. Bioinformatics 28: 423–425.

Carvalho CMB, Lupski JR. 2016. Mechanisms underlying structural variant formation in genomic disorders. Nat Rev Genet 17: 224–238.

Carvalho CMB, Pehlivan D, Ramocki MB, Fang P, Alleva B, Franco LM, Belmont JW, Hastings PJ, Lupski JR. 2013. Replicative mechanisms for CNV formation are error prone. Nat Genet 45: 1319–1326.

Chiang C, Layer RM, Faust GG, Lindberg MR, Rose DB, Garrison EP, Marth GT, Quinlan AR, Hall IM. 2015. SpeedSeq: ultra-fast personal genome analysis and interpretation. Nat Methods 12: 966–968.

Chong Z, Ruan J, Gao M, Zhou W, Chen T, Fan X, Ding L, Lee AY, Boutros P, Chen J, et al. 2017. novoBreak: local assembly for breakpoint detection in cancer genomes. Nat Methods 14: 65–67.

Cibulskis K, Lawrence MS, Carter SL, Sivachenko A, Jaffe D, Sougnez C, Gabriel S, Meyerson M, Lander ES, Getz G. 2013. Sensitive detection of somatic point mutations in impure and heterogeneous cancer samples. Nat Biotechnol 31: 213–219.

Cingolani P, Platts A, Wang LL, Coon M, Nguyen T, Wang L, Land SJ, Lu X, Ruden DM. 2012. A program for annotating and predicting the effects of single nucleotide polymorphisms, SnpEff: SNPs in the genome of Drosophila melanogaster strain w1118; iso-2; iso-3. Fly 6: 80–92.

Dutta D, Dobson AJ, Houtz PL, Gläßer C, Revah J, Korzelius J, Patel PH, Edgar BA, Buchon N. 2015. Regional Cell-Specific Transcriptome Mapping Reveals Regulatory Complexity in the Adult Drosophila Midgut. Cell Rep 12: 346–358.

Fang LT, Afshar PT, Chhibber A, Mohiyuddin M, Fan Y, Mu JC, Gibeling G, Barr S, Asadi NB, Gerstein MB, et al. 2015. An ensemble approach to accurately detect somatic mutations using SomaticSeq. Genome Biol 16: 197.

Favorov A, Mularoni L, Cope LM, Medvedeva Y, Mironov AA, Makeev VJ, Wheelan SJ. 2012. Exploring massive, genome scale datasets with the GenometriCorr package. PLoS Comput Biol 8: e1002529.

Gadgil R, Barthelemy J, Lewis T, Leffak M. 2017. Replication stalling and DNA microsatellite instability. Biophys Chem 225: 38–48.

Garrison E, Marth G. 2012. Haplotype-based variant detection from short-read sequencing. arXiv [q-bioGN]. http://arxiv.org/abs/1207.3907.

Gel B, Díez-Villanueva A, Serra E, Buschbeck M, Peinado MA, Malinverni R. 2016. regioneR: an R/Bioconductor package for the association analysis of genomic regions based on permutation tests. Bioinformatics 32: 289–291.

Gerstung M, Jolly C, Leshchiner I, Dentro SC, Gonzalez S, Rosebrock D, Mitchell TJ, Rubanova Y, Anur P, Yu K, et al. 2020. The evolutionary history of 2,658 cancers. Nature 578: 122–128.

Gervais L, van den Beek M, Josserand M, Sallé J, Stefanutti M, Perdigoto CN, Skorski P, Mazouni K, Marshall OJ, Brand AH, et al. 2019. Stem Cell Proliferation Is Kept in Check by the Chromatin Regulators Kismet/CHD7/CHD8 and Trr/MLL3/4. Dev Cell 49: 556–573.e6.

Glodzik D, Morganella S, Davies H, Simpson PT, Li Y, Zou X, Diez-Perez J, Staaf J, Alexandrov LB, Smid M, et al. 2017. A somatic-mutational process recurrently duplicates germline susceptibility loci and tissue-specific super-enhancers in breast cancers. Nat Genet 49: 341–348.

Grant CE, Bailey TL, Noble WS. 2011. FIMO: scanning for occurrences of a given motif. Bioinformatics 27: 1017–1018.

Greenman C, Stephens P, Smith R, Dalgliesh GL, Hunter C, Bignell G, Davies H, Teague J, Butler A, Stevens C, et al. 2007. Patterns of somatic mutation in human cancer genomes. Nature 446: 153–158.

Hastings PJ, Ira G, Lupski JR. 2009. A microhomology-mediated break-induced replication model for the origin of human copy number variation. PLoS Genet 5: e1000327.

Huang X, Madan A. 1999. CAP3: A DNA sequence assembly program. Genome Res 9: 868–877.

ICGC/TCGA Pan-Cancer Analysis of Whole Genomes Consortium. 2020. Pan-cancer analysis of whole genomes. Nature 578: 82–93.

Katainen R, Dave K, Pitkänen E, Palin K, Kivioja T, Välimäki N, Gylfe AE, Ristolainen H, Hänninen UA, Cajuso T, et al. 2015. CTCF/cohesin-binding sites are frequently mutated in cancer. Nat Genet 47: 818–821.

Keller I, Bensasson D, Nichols RA. 2007. Transition-transversion bias is not universal: a counter example from grasshopper pseudogenes. PLoS Genet 3: e22.

Kidd JM, Graves T, Newman TL, Fulton R, Hayden HS, Malig M, Kallicki J, Kaul R, Wilson RK, Eichler EE. 2010. A human genome structural variation sequencing resource reveals insights into mutational mechanisms. Cell 143: 837–847.

Kim S, Scheffler K, Halpern AL, Bekritsky MA, Noh E, Källberg M, Chen X, Kim Y, Beyter D, Krusche P, et al. 2018. Strelka2: fast and accurate calling of germline and somatic variants. Nat Methods 15: 591–594.

Koboldt DC, Zhang Q, Larson DE, Shen D, McLellan MD, Lin L, Miller CA, Mardis ER, Ding L, Wilson RK. 2012. VarScan 2: somatic mutation and copy number alteration discovery in cancer by exome sequencing. Genome Res 22: 568–576.

Kurahashi H, Inagaki H, Yamada K, Ohye T, Taniguchi M, Emanuel BS, Toda T. 2004. Cruciform DNA structure underlies the etiology for palindrome-mediated human chromosomal translocations. J Biol Chem 279: 35377–35383.

Larson DE, Harris CC, Chen K, Koboldt DC, Abbott TE, Dooling DJ, Ley TJ, Mardis ER, Wilson RK, Ding L. 2012. SomaticSniper: identification of somatic point mutations in whole genome sequencing data. Bioinformatics 28: 311–317.

Layer RM, Chiang C, Quinlan AR, Hall IM. 2014. LUMPY: a probabilistic framework for structural variant discovery. Genome Biol 15: R84.

Lee JA, Carvalho CMB, Lupski JR. 2007. A DNA replication mechanism for generating nonrecurrent rearrangements associated with genomic disorders. Cell 131: 1235–1247.

Lee-Six H, Olafsson S, Ellis P, Osborne RJ, Sanders MA, Moore L, Georgakopoulos N, Torrente F, Noorani A, Goddard M, et al. 2019. The landscape of somatic mutation in normal colorectal epithelial cells. Nature 574: 532–537.

Letouzé E, Shinde J, Renault V, Couchy G, Blanc J-F, Tubacher E, Bayard Q, Bacq D, Meyer V, Semhoun J, et al. 2017. Mutational signatures reveal the dynamic interplay of risk factors and cellular processes during liver tumorigenesis. Nat Commun 8: 1315.

Liu P, Carvalho CMB, Hastings PJ, Lupski JR. 2012. Mechanisms for recurrent and complex human genomic rearrangements. Curr Opin Genet Dev 22: 211–220.

Li Y, Roberts ND, Wala JA, Shapira O, Schumacher SE, Kumar K, Khurana E, Waszak S, Korbel JO, Haber JE, et al. 2020. Patterns of somatic structural variation in human cancer genomes. Nature 578: 112–121.

Lu S, Wang G, Bacolla A, Zhao J, Spitser S, Vasquez KM. 2015. Short Inverted Repeats Are Hotspots for Genetic Instability: Relevance to Cancer Genomes. Cell Rep 10: 1674–1680.

Martincorena I, Fowler JC, Wabik A, Lawson ARJ, Abascal F, Hall MWJ, Cagan A, Murai K, Mahbubani K, Stratton MR, et al. 2018. Somatic mutant clones colonize the human esophagus with age. Science 362: 911–917.

Martincorena I, Raine KM, Gerstung M, Dawson KJ, Haase K, Van Loo P, Davies H, Stratton MR, Campbell PJ. 2017. Universal Patterns of Selection in Cancer and Somatic Tissues. Cell 171: 1029–1041.e21.

Martincorena I, Roshan A, Gerstung M, Ellis P, Van Loo P, McLaren S, Wedge DC, Fullam A, Alexandrov LB, Tubio JM, et al. 2015. High burden and pervasive positive selection of somatic mutations in normal human skin. Science 348: 880–886.

McKenna A, Hanna M, Banks E, Sivachenko A, Cibulskis K, Kernytsky A, Garimella K, Altshuler D, Gabriel S, Daly M, et al. 2010. The Genome Analysis Toolkit: a MapReduce framework for analyzing next-generation DNA sequencing data. Genome Res 20: 1297–1303.

Micchelli CA, Perrimon N. 2006. Evidence that stem cells reside in the adult Drosophila midgut epithelium. Nature 439: 475–479.

modENCODE Consortium, Roy S, Ernst J, Kharchenko PV, Kheradpour P, Negre N, Eaton ML, Landolin JM, Bristow CA, Ma L, et al. 2010. Identification of functional elements and regulatory circuits by Drosophila modENCODE. Science 330: 1787–1797.

Moore L, Leongamornlert D, Coorens THH, Sanders MA, Ellis P, Dentro SC, Dawson KJ, Butler T, Rahbari R, Mitchell TJ, et al. 2020. The mutational landscape of normal human endometrial epithelium. Nature 1–7.

Nazaryan-Petersen L, Eisfeldt J, Pettersson M, Lundin J, Nilsson D, Wincent J, Lieden A, Lovmar L, Ottosson J, Gacic J, et al. 2018. Replicative and non-replicative mechanisms in the formation of clustered CNVs are indicated by whole genome characterization. PLoS Genet 14: e1007780.

Nik-Zainal S, Alexandrov LB, Wedge DC, Van Loo P, Greenman CD, Raine K, Jones D, Hinton J, Marshall J, Stebbings LA, et al. 2012. Mutational processes molding the genomes of 21 breast cancers. Cell 149: 979–993.

Nik-Zainal S, Davies H, Staaf J, Ramakrishna M, Glodzik D, Zou X, Martincorena I, Alexandrov LB, Martin S, Wedge DC, et al. 2016. Landscape of somatic mutations in 560 breast cancer whole-genome sequences. Nature 534: 47–54.

Ohlstein B, Spradling A. 2006. The adult Drosophila posterior midgut is maintained by pluripotent stem cells. Nature 439: 470–474.

Ostrow SL, Barshir R, DeGregori J, Yeger-Lotem E, Hershberg R. 2014. Cancer evolution is associated with pervasive positive selection on globally expressed genes. PLoS Genet 10: e1004239.

Paeschke K, Capra JA, Zakian VA. 2011. DNA replication through G-quadruplex motifs is promoted by the Saccharomyces cerevisiae Pif1 DNA helicase. Cell 145: 678–691.

Palmer WH, Medd NC, Beard PM, Obbard DJ. 2018. Isolation of a natural DNA virus of Drosophila melanogaster, and characterisation of host resistance and immune responses. PLoS Pathog 14: e1007050.

Petrov DA, Hartl DL. 1999. Patterns of nucleotide substitution in Drosophila and mammalian genomes. Proc Natl Acad Sci U S A 96: 1475–1479.

Pleasance ED, Cheetham RK, Stephens PJ, McBride DJ, Humphray SJ, Greenman CD, Varela I, Lin M-L, Ordóñez GR, Bignell GR, et al. 2010. A comprehensive catalogue of somatic mutations from a human cancer genome. Nature 463: 191–196.

Quinlan AR, Hall IM. 2010. BEDTools: a flexible suite of utilities for comparing genomic features. Bioinformatics 26: 841–842.

Raddatz G, Guzzardo PM, Olova N, Fantappié MR, Rampp M, Schaefer M, Reik W, Hannon GJ, Lyko F. 2013. Dnmt2-dependent methylomes lack defined DNA methylation patterns. Proc Natl Acad Sci U S A 110: 8627–8631.

Rausch T, Zichner T, Schlattl A, Stütz AM, Benes V, Korbel JO. 2012. DELLY: structural variant discovery by integrated paired-end and split-read analysis. Bioinformatics 28: i333–i339.

Rheinbay E, Nielsen MM, Abascal F, Wala JA, Shapira O, Tiao G, Hornshøj H, Hess JM, Juul RI, Lin Z, et al. 2020. Analyses of non-coding somatic drivers in 2,658 cancer whole genomes. Nature 578: 102–111.

Robberecht C, Voet T, Zamani Esteki M, Nowakowska BA, Vermeesch JR. 2013. Nonallelic homologous recombination between retrotransposable elements is a driver of de novo unbalanced translocations. Genome Res 23: 411–418.

Schuster-Böckler B, Lehner B. 2012. Chromatin organization is a major influence on regional mutation rates in human cancer cells. Nature 488: 504–507.

Siudeja K, Nassari S, Gervais L, Skorski P, Lameiras S, Stolfa D, Zande M, Bernard V, Frio TR, Bardin AJ. 2015. Frequent Somatic Mutation in Adult Intestinal Stem Cells Drives Neoplasia and Genetic Mosaicism during Aging. Cell Stem Cell 17: 663–674.

Siudeja K, van den Beek M, Riddiford N, Boumard B, Wurmser A, Stefanutti M, Bardin A. 2020. Somatic transposition in the fly intestine. bioRxiv (deposition ongoing).

Supek F, Miñana B, Valcárcel J, Gabaldón T, Lehner B. 2014. Synonymous mutations frequently act as driver mutations in human cancers. Cell 156: 1324–1335.

Tubbs A, Sridharan S, van Wietmarschen N, Maman Y, Callen E, Stanlie A, Wu W, Wu X, Day A, Wong N, et al. 2018. Dual Roles of Poly(dA:dT) Tracts in Replication Initiation and Fork Collapse. Cell 174: 1127–1142.e19.

Woo YH, Li W-H. 2012. DNA replication timing and selection shape the landscape of nucleotide variation in cancer genomes. Nat Commun 3: 1004.

Xie C, Tammi MT. 2009. CNV-seq, a new method to detect copy number variation using high-throughput sequencing. BMC Bioinformatics 10: 80.

Yang L, Luquette LJ, Gehlenborg N, Xi R, Haseley PS, Hsieh C-H, Zhang C, Ren X, Protopopov A, Chin L, et al. 2013. Diverse mechanisms of somatic structural variations in human cancer genomes. Cell 153: 919–929.

Yokoyama A, Kakiuchi N, Yoshizato T, Nannya Y, Suzuki H, Takeuchi Y, Shiozawa Y, Sato Y, Aoki K, Kim SK, et al. 2019. Age-related remodelling of oesophageal epithelia by mutated cancer drivers. Nature 565: 312–317.

Zou X, Morganella S, Glodzik D, Davies H, Li Y, Stratton MR, Nik-Zainal S. 2017. Short inverted repeats contribute to localized mutability in human somatic cells. Nucleic Acids Res. http://academic.oup.com/nar/article/doi/10.1093/nar/gkx731/4090957/Short-inverted-repeats-contribute-to-localized (Accessed October 11,2017).

