## Supplementary Figures and Methods for "Evolution and genomic signatures of spontaneous somatic mutation in *Drosophila* intestinal stem cells"

### Fig. S1

a

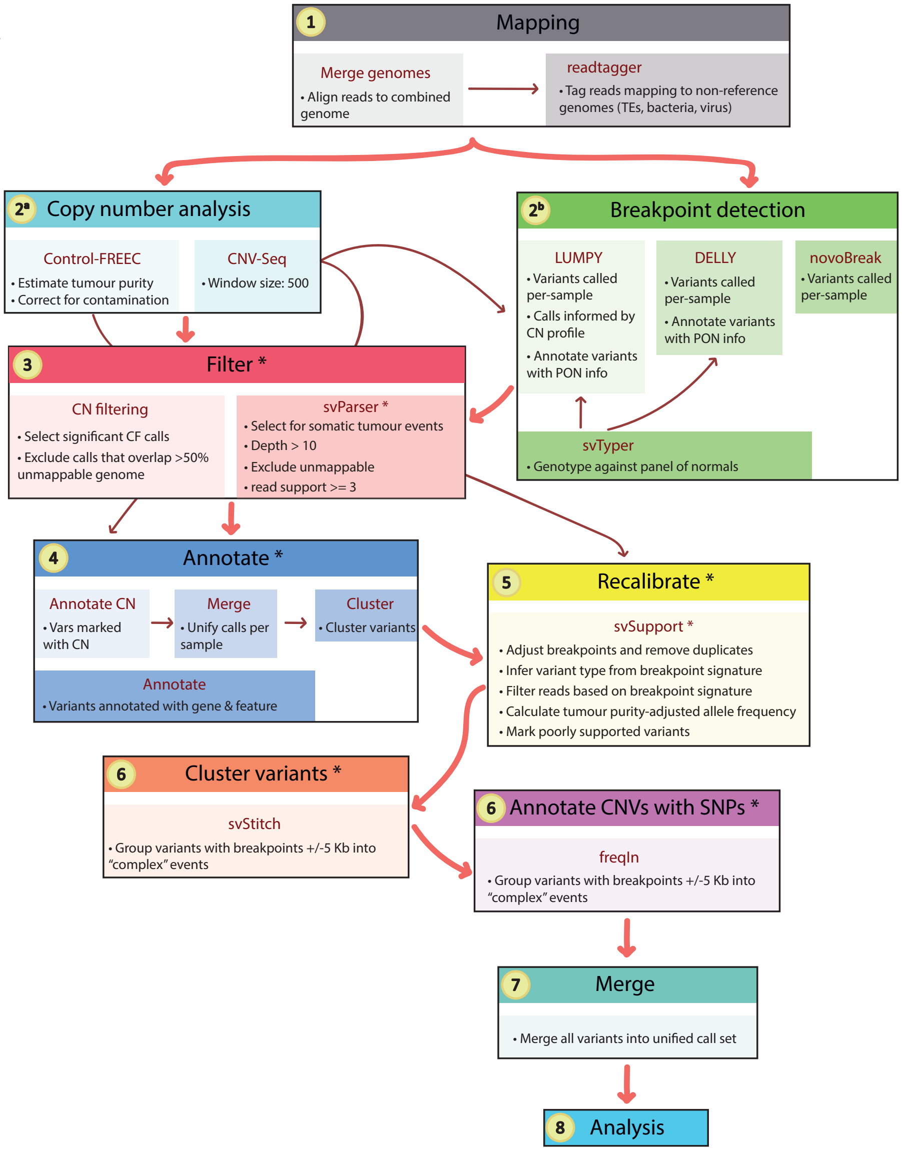

1 Mapping
Merge genomes
• Align reads to combined genome
readtagger
• Tag reads mapping to non-reference genomes (TEs, bacteria, virus)
2a Copy number analysis
Control-FREEC
• Estimate tumour purity
• Correct for contamination
CNV-Seq
• Window size: 500
2b Breakpoint detection
LUMPY
• Variants called per-sample
• Calls informed by CN profile
• Annotate variants with PON info
DELLY
• Variants called per-sample
• Annotate variants with PON info
novoBreak
• Variants called per-sample
svTyper
• Genotype against panel of normals
3 Filter *
CN filtering
• Select significant CF calls
• Exclude calls that overlap >50% unmappable genome
svParser *
• Select for somatic tumour events
• Depth > 10
• Exclude unmappable
• read support >= 3
4 Annotate *
Annotate CN
• Vars marked with CN
Merge
• Unify calls per sample
Cluster
• Cluster variants
Annotate
• Variants annotated with gene & feature
5 Recalibrate *
svSupport *
• Adjust breakpoints and remove duplicates
• Infer variant type from breakpoint signature
• Filter reads based on breakpoint signature
• Calculate tumour purity-adjusted allele frequency
• Mark poorly supported variants
6 Cluster variants *
svStitch
• Group variants with breakpoints +/-5 Kb into "complex" events
6 Annotate CNVs with SNPs *
freqIn
• Group variants with breakpoints +/-5 Kb into "complex" events
7 Merge
• Merge all variants into unified call set
8 Analysis


b

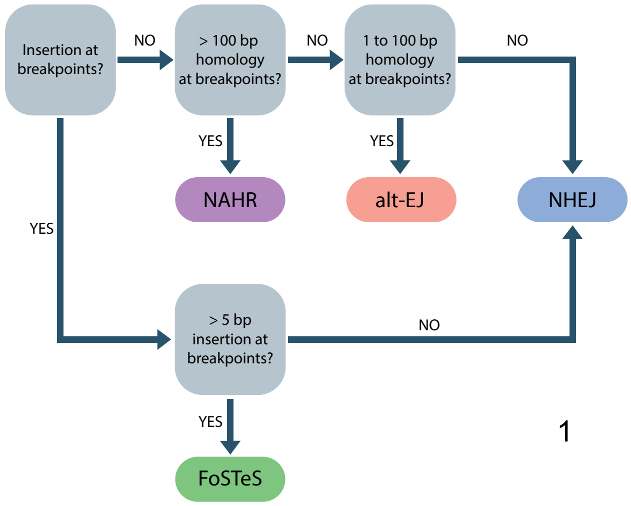

Insertion at breakpoints?
NO
> 100 bp homology at breakpoints?
NO
1 to 100 bp homology at breakpoints?
NO
YES
NAHR
YES
alt-EJ
NHEJ
YES
> 5 bp insertion at breakpoints?
NO
YES
FoSTeS

**Fig. S1. Schematic showing a bioinformatic pipeline for identifying structural variants** (**a**) An overview of the main steps involved in our bioinformatic pipeline. We first map reads to *Drosophila* transposable element sequences as well as multiple genomes found as contaminants in our sequencing data (Methods; Supplementary Methods). Structural variant discovery was then performed in two complementary but distinct steps. First, read depth-based approaches (CNV-Seq (Xie and Tammi 2009); Control-FREEC (Boeva et al. 2012)) were used to detect copy number variants (CNVs). Secondly, we used read mapping-based approaches, utilising aberrantly mapped and/or split-reads (LUMPY (Layer et al. 2014); DELLY (Rausch et al. 2012); novobreak (Chong et al. 2017)) to detect precise breakpoints of multiple classes of structural variant (Fig. 1b; Fig. S1a; Methods). To ensure only true somatic events were considered, a panel of normals (PON) was constructed by combining variant calls from all sequenced normal samples, and used to select for somatic (tumour-only) calls. We then combined variants into unified per-sample call sets, before merging variants that were found by multiple approaches, and then annotated calls with the location of the breakpoints with respect to gene features (Methods). Finally, breakpoints within close proximity (+/- 5 kb) were clustered into “complex” events, and CNVs were annotated with the frequency of heterozygous SNPs, in order to discern false positives (Methods; Supplementary Methods). (**b**) Assigning putative underlying mechanisms of structural variants by analysis of breakpoint junctions. We used annotated breakpoints for microhomology and inserted sequences, and classified each breakpoint junction using criteria from (Yang et al. 2013; Kidd et al. 2010).

### Fig. S2

a

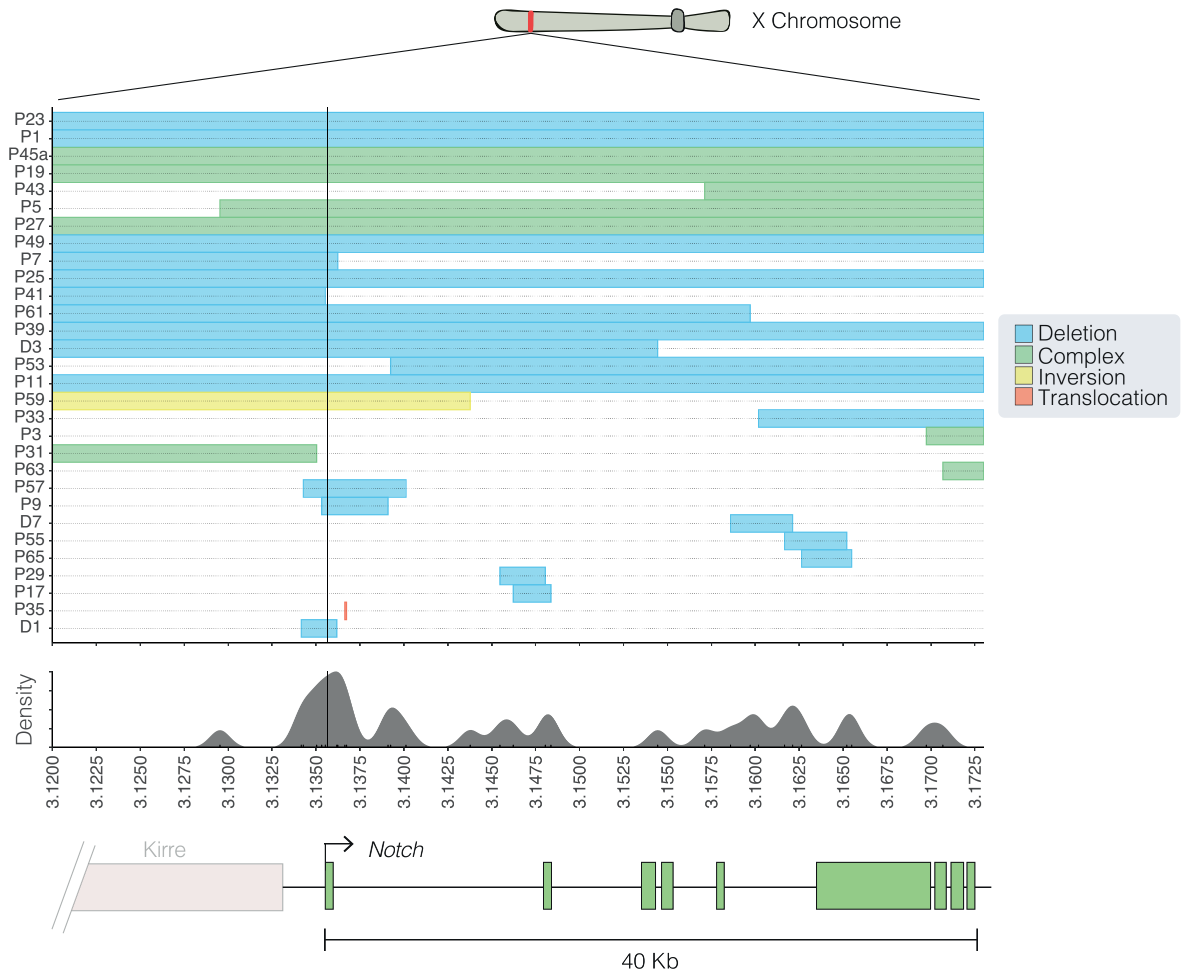


**Fig. S2. The distribution of structural variant breakpoints over the *Notch* locus**
Breakpoints did not occur at the same genomic locus, but clustering (density plot) of breakpoints was found around the TSS of *Notch* (black vertical line).

Fig. S3

a

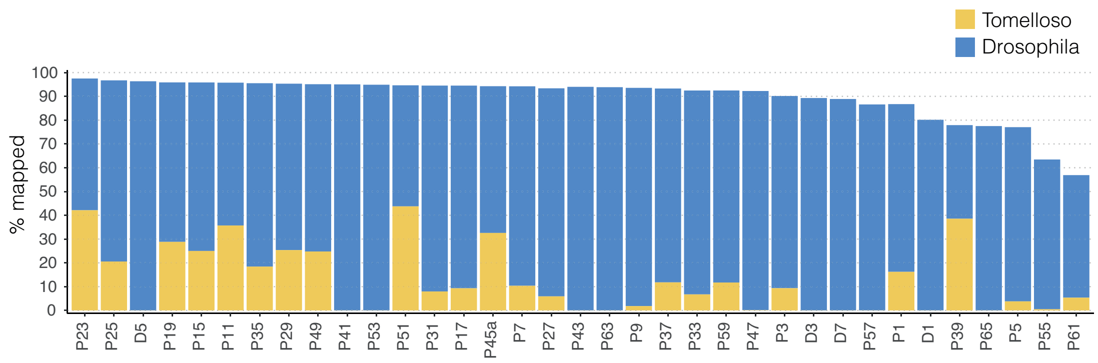


b

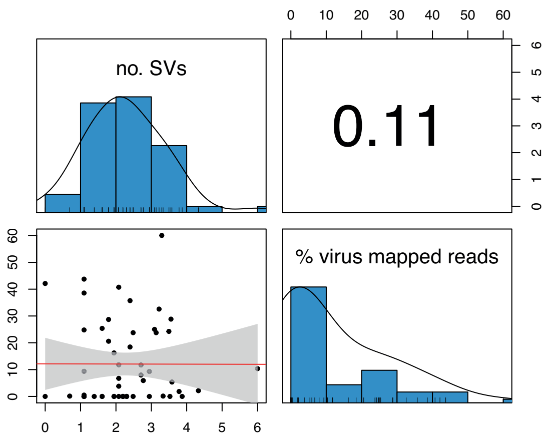


**Fig. S3. Viral contamination is not correlated with mutation count**
(**a**) Percentage of reads mapping to the virus Tomelloso and *Drosophila* genome per tumour sample. (**b**) Pearson's correlation between the number of mutations detected per sample and the percentage of reads mapping to the Tomelloso genome.

### Fig. S4

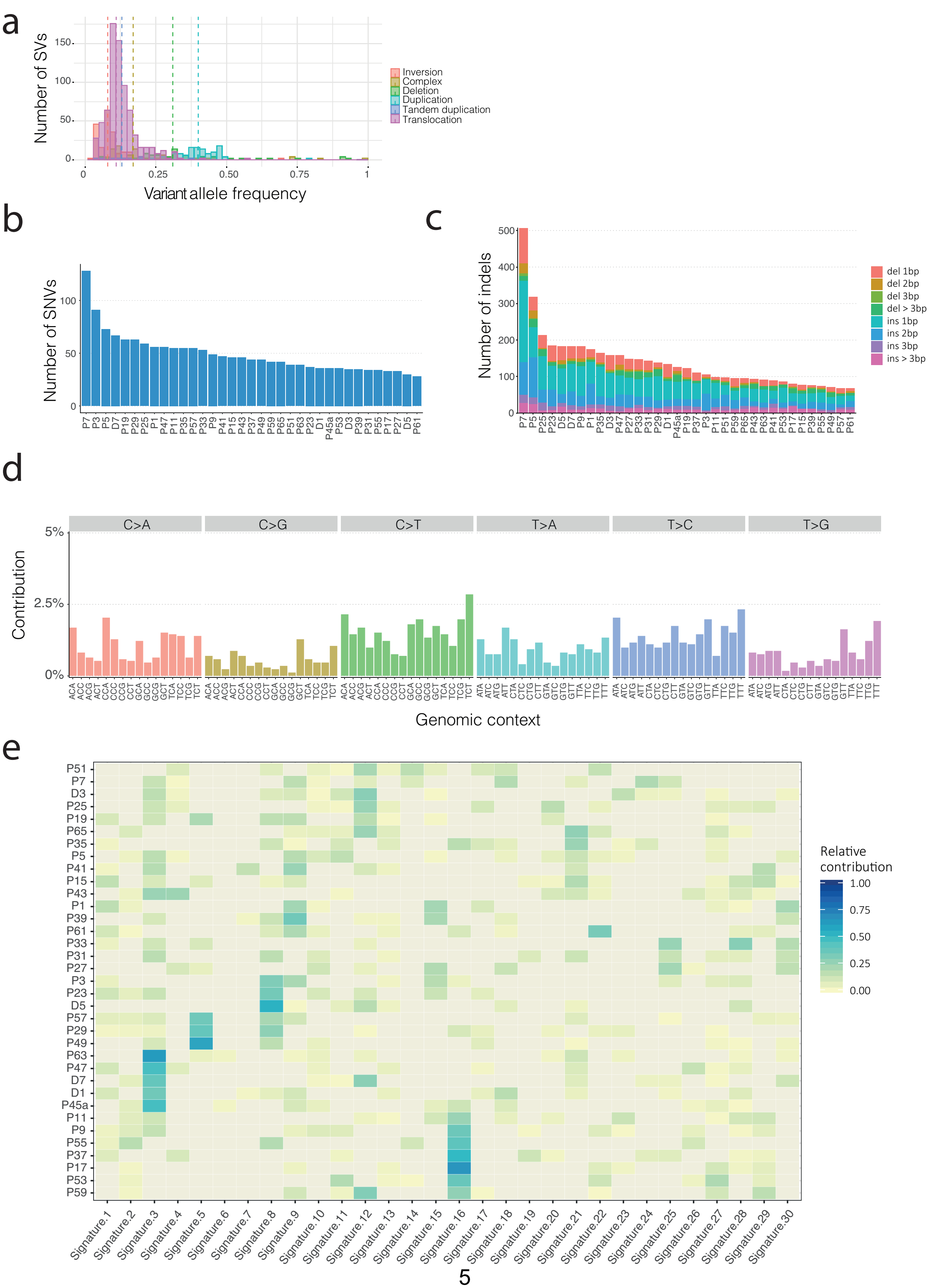
a
Number of SVs
Variant allele frequency
Inversion
Complex
Deletion
Duplication
Tandem duplication
Translocation
b
Number of SNVs
c
Number of indels
del 1bp
del 2bp
del 3bp
del > 3bp
ins 1bp
ins 2bp
ins 3bp
ins > 3bp
d
Contribution
C>A
C>G
C>T
T>A
T>C
T>G
Genomic context
e
Relative contribution

**Fig. S4. Genome-wide mutations**

(**a**) The distribution of variant allele frequency of structural variants observed genome-wide, coloured by class. Number of SNVs (**b**) and indels (**c**) detected genome-wide. (**d**) Distribution of SNVs within a trinucleotide context across samples. (**e**) Per-sample cosine similarity (relative contribution) between observed mutational spectra and COSMIC mutational signatures.

### Fig. S5

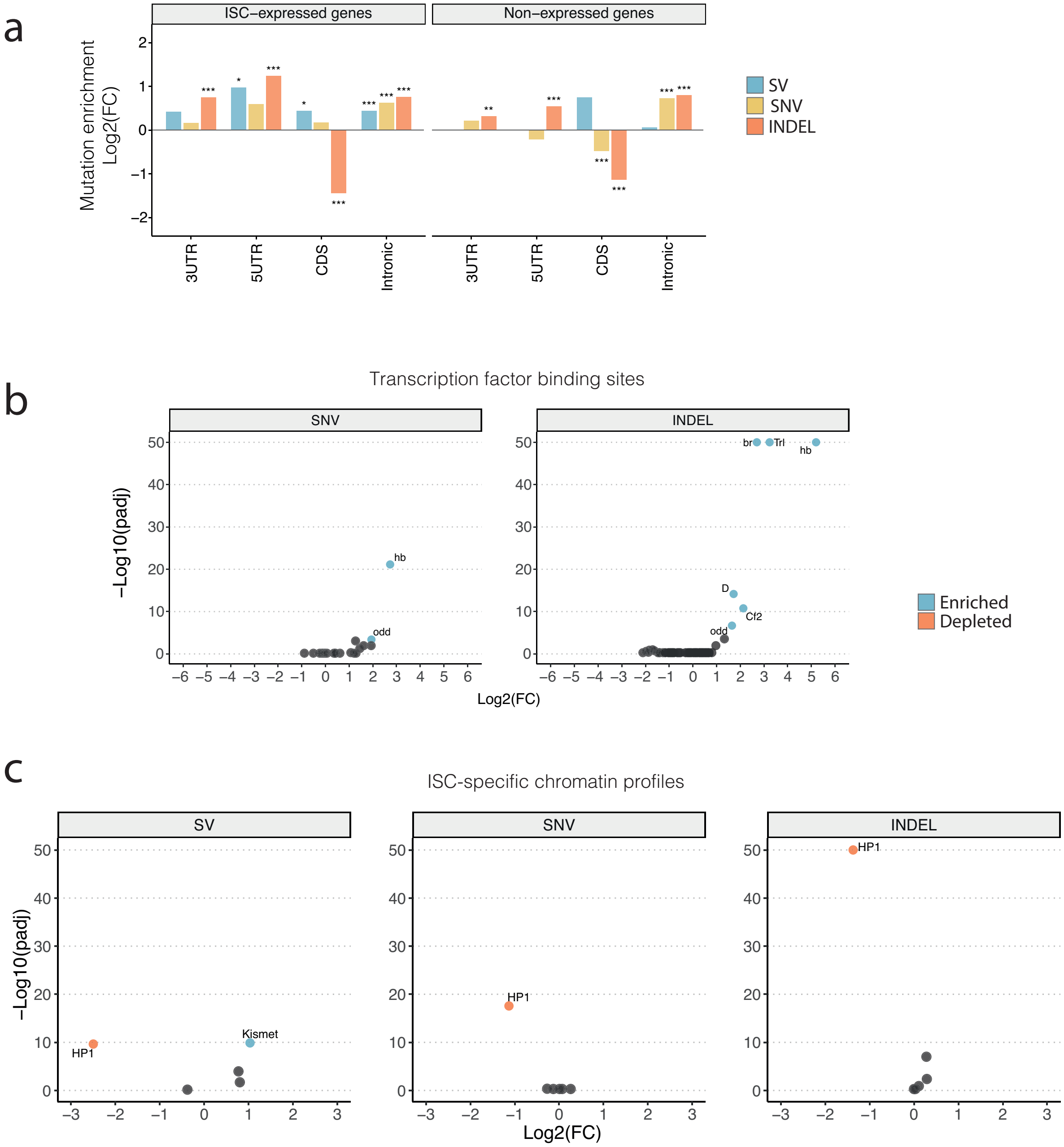
a
ISC–expressed genes
Non–expressed genes
Mutation enrichment
Log2(FC)
SV
SNV
INDEL
3UTR
5UTR
CDS
Intronic
b
Transcription factor binding sites
SNV
INDEL
–Log10(padj)
Log2(FC)
hb
odd
br
Trl
D
Cf2
Enriched
Depleted
c
ISC-specific chromatin profiles
SV
SNV
INDEL
HP1
Kismet

**Fig. S5. Distribution of somatic mutations in genome features**
(**a**) Enrichment of mutations in ISC-expressed vs non-expressed genes. Indels were strongly depleted in CDS regions. (**b**, **c**) Volcano plots showing enrichment or depletion of mutations in TF binding sites (**b**) and chromatin landscape (**c**). Highlighted features represent those that with an E-score (-Log10(p)*Log2(FC); Methods) > 5. The y axis of **b** and **c** are restricted to a maximum -Log10(padj) value of 50 Asterisks denote significance: *** $p < 0.001$; ** $p < 0.01$; * $p < 0.5$. All p values shown have been generated from a two-sided binomial test, and adjusted for multiple comparisons using a Benjamini-Hochberg adjustment.

Fig. S6

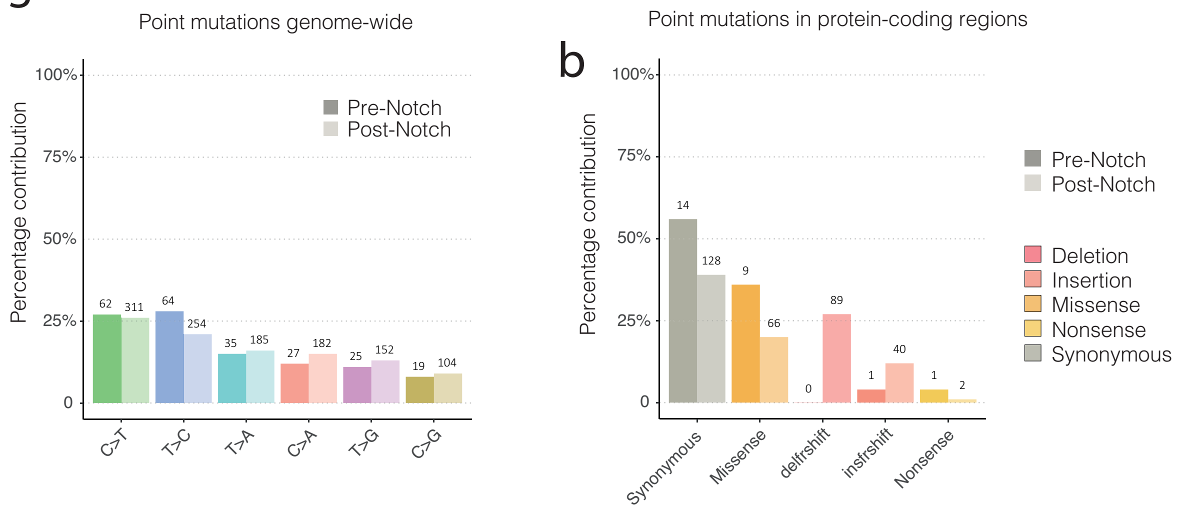


**Fig. S6. Observed mutational frequencies occurring pre- and post-*Notch* inactivation**
(**a**) Frequency of nucleotide substitution types were consistent between groups. (**b**) Frequency of protein-coding point mutation types. We observed no small frame-shift-inducing deletions occurring pre-*Notch* inactivation.

#### Supplementary Methods

##### Creating a panel of normals (PON) with Mutect2 (v4.1.2)

For each normal sample we ran Mutect2 with the `--disable-read-filter MateOnSameContigOrNoMappedMateReadFilter` option.

##### Mutect2 (v4.1.2)

For each tumour normal pair we ran Mutect2 specifying the unmappable regions of the genome as `--exclude-intervals`, and we used the `--dont-use-soft-clipped-bases` option to remove SNV calling in noisy reads. We specified the PON with `--panel-of-normals`. We then further filtered calls using the `FilterMutectCalls` function.

##### Varscan2 (v2.4)

For each tumour normal pair, we ran Varscan2 in somatic mode from combined pileups generated from both bam files using `--mpileup 1`. We specified minimum coverage of 20 for both the tumour and normal sample using `--min-coverage-normal 20 --min-coverage-tumor 20` and provided the tumour purity value estimated from Control-FREEC using `--tumor-purity`. We applied a strand filter using `--strand-filter 1`. We then ran `processSomatic` to separate high-confidence somatic SNV and indel calls. Next, we excluded high-quality somatic calls within the unmappable genome and removed sites that were also found in the PON.

##### Strelka (v2.9.10)

For each tumour normal pair we ran `configureStrelkaSomaticWorkflow.py` followed by `runWorkflow.py -m local -j 4 -g 5`.

**SomaticSniper (v1.0.5.0)**

For each tumour normal pair we ran `bam-somaticsniper -Q 40 -L`.

**SomaticSeq (v3.3.0)**

To combine calls made by multiple callers, for each tumour normal pair we ran `somaticseq_parallel.py --algorithm ada` specifying the mappable genome as ` --inclusion-region`.

**Freebayes (v1.2.0-dirty)**

For each tumour normal pair we ran `freebayes -0  --pooled-discrete --genotype-qualities`, specifying a minimum coverage of 20 with `--min-coverage 20`. Next, we excluded calls within the unmappable genome, and processed calls using the vcflib function `vcfallelicprimitives` (https://github.com/vcflib/vcflib) and vt (https://github.com/atks/vt) using `vt decompose_blocksub` to decompose biallelic block substitutions into their constituent SNVs. We then normalised calls using `vt normalize -q`, and selected somatic calls using the vcflib function `vcfsamplediff`. We then filtered somatic calls using the vcflib function `vcffilter` to select for those with a depth greater than 20 using `vcffilter -f "DP > 20"`, for high-quality calls `vcffilter -f "QUAL > 1 & QUAL / AO > 10"` and for those supported by reads on both DNA strands `vcffilter -f "SAF > 0 & SAR > 0"` and for those with both right- and left-facing read support `vcffilter -f "RPR > 0 & RPL > 0"`.

**Combining point mutations**

We then merged the somatic calls from Freebayes with the output of SomaticSeq and re-filtered the merged per-sample calls against the PON using https://github.com/bardin-lab/mutationProfiles/blob/master/script/filtervcf.py.

**Annotating point mutations and downstream analysis**

Merged and filtered point mutations were annotated for gene features and functional impact estimates using snpEff v4.3. We then used the R package dNdScv (Martincorena et al. 2017) to annotate protein-coding mutations for their functional impact.

**Downstream analysis**

Downstream analyses were performed using functions developed within the mutationProfiles suite of tools (https://github.com/bardin-lab/mutationProfiles) and are detailed in our pipeline (https://github.com/bardin-lab/Riddiford_et_al_2020/blob/master/sv_analysis.R).

**CNV-Seq**

We first created a file containing read counts for each bam file using `samtools view ${bam_file} | perl -lane 'print "$F[2]\t$F[3]"' > ${out}.hits`. We then ran CNV-Seq on tumour normal pairs using window sizes of 500 bps and 50 kb with `cnv-seq.pl --window-size $window --genome-size 137547960 --global-normalization`. For small window sizes we then filtered windows with fewer than 50 reads in the normal sample.

**Control-FREEC (v11.0)**

For each tumour normal pair we ran Control-FREEC as described in the documentation, and annotated significant CNVs using `assess_significance.R`. We then filtered for significant CNVs, and wrote .gff3 files for inspection in IGV. We then further filtered CNV calls to exclude those that overlapped more than 25% with unmappable regions of the genome using bedtools subtract v2.28.0 (Quinlan and Hall 2010).

**novoBreak (v1.1)**

We ran novoBreak for each tumour normal pair using the recommended filtering steps.

**LUMPY (v0.2.13)**

For each tumour normal pair we first extracted discordant and split reads and estimated the insert size distribution as described (https://github.com/arq5x/lumpy-sv). We then ran LUMPY using paired-end and split-reads and excluding the unmappable genome using `-x`.

**svTyper (svtools 0.3.0)**

We then created a panel of normals (PON) using svTyper run on all normal samples and used this to genotype calls made for each tumour normal pair.

**DELLY (v0.7.8)**

For each tumour normal pair we ran `delly call`, excluding unmappable regions using the `-x` option. We then ran `delly filter -f somatic` to select for somatic calls and then genotyped calls against the PON generated by svTools by running `delly call` on the output generated in the above step. We then post-filtered for somatic calls after genotyping.

**Structural variant filtering and annotation**

In order to filter, annotate and combine calls made from the different approaches described above, we developed a suite of tools, svParser (https://github.com/bardin-lab/svParser). By inspecting the evidence behind individual calls using `perl script/svParse.pl -i`, and viewing evidence in IGV, we optimised a set of filters that appeared to be false positives while retaining calls that were well supported. We developed a wrapper script to run our pipeline `runParser -fmas`, that runs `svParse` on per-sample variant calls made by LUMPY, novoBreak and DELLY with the filters `-f chr=1 -f su=3 -f dp=10 -f sq=0.1 -f rdr=0.05 -s`. We use the `-c` option to specify a directory containing CNV-Seq count files for each tumour normal pair, which we use to annotate average Log2(FC) values over CNV regions. After annotating breakpoints in genes for the gene name and gene feature we inspected variants in IGV, and excluded several events that appeared to be false positive duplications called due to inconsistent coverage between the tumour and normal samples that was not consistent with a duplication in the tumour sample (Supplementary Table 5).

**Re-calibrating structural variant breakpoints**

In order to standardise calls made between different approaches, and refine breakpoint calls, we developed svSupport (https://github.com/bardin-lab/svSupport). First, for each variant with split-read support, we extracted a breakpoint signature to classify events as deletions, tandem duplications, inversions or translocations. For each breakpoint, we search for reads that are tagged as mapping to a non-*Drosophila* chromosome, and in filter variants with inconsistent support between breakpoints, or where we can't find any split-read support after duplicate removal. We also annotate breakpoints that are associated with TE-tagged reads with the number of supporting reads and the TE class. Variants with low read support (< 3 split-reads) were recorded and later removed. We then incorporated tumour purity values estimated by Control-FREEC to adjust the allele frequency of each variant. Here, we first extracted the number of reads directly supporting or opposing a given breakpoint and then calculated an adjusted opposing read count given the purity of the sample as follows: `expected_oppose = (1 - tumour_purity) * total_reads`. We then use this adjusted opposing read counts to calculate a tumour purity adjusted allele frequency. For variants with imprecise breakpoints (CN events called by read-depth-based approaches) we first counted the number of reads mapped in the CNV region in both the tumour and normal sample. Next we normalised read counts for sequencing depth by counting the number of reads mapped across all full chromosomes (2L, 2R, 3L, 3R, 4, X and Y). We then performed a similar adjustment as described above for split-read-supported variants to adjust allele frequency. Here, the supporting reads were derived by subtracting the read count in the tumour sample from the read count in the normal sample. The opposing reads were then calculated by subtracting the supporting read count from the normal read count, and a tumour purity-adjusted allele frequency calculated as above.

**Clustering breakpoints**

In order to identify complex events from clusters of linked breakpoints, we used a tool developed in the svParser suite (https://github.com/bardin-lab/svParser/blob/master/script/svStitch.py). For each sample we search for variants whose breakpoints were within a 5 kb window. In cases where clustered variants all belonged to the same class of CN event (deletion or duplication) individual variants were collapsed into a single event and the variant type was not modified. In cases where multiple classes of structural variant class were clustered, or all classes belonged to the same class but were not CN events, we re-annotated each variant type in the cluster as "complex", and collapsed variants into a single mutational event.

**Annotating CNVs with SNP frequencies**

As a final filtering step, we used germline SNP calls made by Freebayes to remove spurious CNV calls. First, allele frequencies of heterozygous germline SNPs called at sites with a depth > 20 in both the tumour and normal sample were extracted over a CN region. For each SNP, we calculated the difference in frequency between the tumour and normal sample, and recorded the difference as CNV-supporting if it was > 10%. We recorded the number of informative SNPs over CNVs, and in cases where we had sufficient evidence to reject a CN call (> 5 informative SNPs and supporting SNPs / opposing SNPs < 2) we marked variants as false positives.

**Downstream analysis**

Downstream analyses were performed using functions developed within the svBreaks suite of tools (https://github.com/bardin-lab/svBreaks), and are detailed in our pipeline (https://github.com/bardin-lab/Riddiford_et_al_2020/blob/master/sv_analysis.R).
